# Motor patterning, ion regulation and Spreading Depolarization during CNS shutdown induced by experimental anoxia in *Locusta migratoria*

**DOI:** 10.1101/2021.05.12.443850

**Authors:** R. Meldrum Robertson, Rachel A. Van Dusen

## Abstract

Anoxia induces a reversible coma in insects. Coma onset is triggered by the arrest of mechanisms responsible for maintaining membrane ion homeostasis in the CNS, resulting in a wave of neuronal and glial depolarization known as spreading depolarization (SD). Different methods of anoxia influence the behavioural response but their effects on SD are unknown. We investigated the effects of CO_2_, N_2_, and H_2_O on the characteristics of coma induction and recovery in *Locusta migratoria*. Water immersion delayed coma onset and recovery, likely due to involvement of the tracheal system and the nature of asphyxiation but otherwise resembled N_2_ delivery. The main difference between N_2_ and CO_2_ was that CO_2_ hastened onset of neural failure and SD and delayed recovery. In the CNS, this was associated with CO_2_ inducing an abrupt and immediate decrease of interstitial pH and increase of extracellular [K^+^]. Recording of the transperineurial potential showed that SD propagation and a postanoxic negativity (PAN) were similar with both gases. The PAN increased with ouabain treatment, likely due to removal of the counteracting electrogenic effect of Na^+^/K^+^-ATPase, and was inhibited by bafilomycin, a proton pump inhibitor, suggesting that it was generated by the electrogenic effect of a Vacuolar-type ATPase (VA). Muscle fibres depolarized by ∼20 mV, which happened more rapidly with CO_2_ compared with N_2_. Wing muscle motoneurons depolarized nearly completely in two stages, with CO_2_ causing more rapid onset and slower recovery than N_2_. Other parameters of SD onset and recovery were similar with the two gases. Electrical resistance across the ganglion sheath increased during anoxia and at SD onset. We provisionally attribute this to cell swelling reducing the dimensions of the interstitial pathway from neuropil to the bathing saline. Neuronal membrane resistance decreased abruptly at SD onset indicating opening of an unidentified membrane conductance. Consideration of the intracellular recording relative to the saline suggests that the apical membrane of perineurial glia depolarizes prior to neuron depolarization. We propose that SD is triggered by events at the perineurial sheath and then propagates laterally and more deeply into the neuropil. We conclude that the fundamental nature of SD is not dependent on the method of anoxia however the timing of onset and recovery are influenced; water immersion is complicated by the tracheal system and CO_2_ delivery has more rapid and longer lasting effects, associated with severe interstitial acidosis.

## Introduction

Hypoxia-tolerant animals achieve their remarkable success by virtue of mechanisms of metabolic arrest that conserve energy (Hochachka, 1986a; Hochachka, 1986b). Given the high energetic cost of information processing in the CNS (Attwell and Laughlin, 2001) a primary target for conservation is CNS ion homeostasis, the costs of which can be greatly reduced by channel arrest and spike arrest (Jonz et al., 2016; Robertson, 2017). Insects can be exposed to severe hypoxia or anoxia in a wide range of habitats (Hoback, 2012; Hoback and Stanley, 2001) and they employ a CNS arrest strategy of metabolic depression via complete neuromuscular shutdown associated with a stress-induced coma, which allows them to delay cellular energy depletion (Campbell et al., 2018; Campbell et al., 2019; Robertson et al., 2017). Although genetic approaches have identified potential molecular mechanisms underlying anoxic coma (Ma et al., 2001; Xiao and Robertson, 2016; Xiao and Robertson, 2017) and shown them to be under selection pressure (Xiao et al., 2019), we do not completely understand what processes determine the resistance to hypoxia or the speed of recovery from the coma. In addition, to expedite future research, it is important to determine the most biologically relevant methodology for inducing experimental anoxia.

Coma in insects and mammals are similar behaviourally (immobility and loss of responsiveness) but the neural mechanisms are different. Coma induction in insects follows a loss of membrane ion homeostasis within the nervous system, leading to a silencing of neural and muscular activity. This results in a large redistribution of ions across neuronal and glial membranes generating a wave of cellular depolarization: a phenomenon known as spreading depolarization (SD) (Dreier and Reiffurth, 2015; Pietrobon and Moskowitz, 2014; Rodgers et al., 2010), which exhibits several characteristics that are similar in both insects and mammals (Robertson et al., 2020; Spong et al., 2016a; Spong et al., 2017). There is considerable clinical interest in SD due to its involvement in several human pathologies including migraine, stroke, and traumatic brain injury (Shuttleworth et al., 2019) and the suggestion that SD may have beneficial effects to terminate and prevent seizure activity (Tamim et al., 2021). In spite of many years of research, there is limited consensus on the mechanisms of SD (Andrew et al., 2021) and insect model systems afford an opportunity to investigate SD without the difficulties associated with mammalian preparations (Spong et al., 2017).

In insects, there are different methods of creating anoxic conditions, which affect the characteristics of the ensuing coma, providing a means to experimentally probe the mechanisms of neural shutdown. Anoxic coma can be induced via asphyxiation, caused by exposure to any gas with less than 2% oxygen or by submersion in water. Carbon dioxide (CO_2_) and nitrogen gas (N_2_) are commonly used to immobilize insects during experimental work, however these gases, particularly CO_2_, can affect subsequent animal behaviour and physiology (Colinet and Renault, 2012; MacMillan et al., 2017; Milton and Partridge, 2008; Perron et al., 1972; Woodring et al., 1978). Water immersion is a common, ecologically relevant cause of anoxia for many insects (Brust et al., 2005; Brust et al., 2007; Plum, 2005; Woodman, 2013; Woodman, 2015) and it has been used to investigate physiological mechanisms of anoxic coma (Benasayag-Meszaros et al., 2015; Hou et al., 2014; Robertson et al., 2019). However, whether water immersion, N_2_ or CO_2_ exposure have different physiological consequences in the CNS is unknown.

We were interested in the mechanisms of CNS shutdown that enhance metabolic depression for short durations (≤ 30 mins) rather than the tissue injury that occurs after long durations of anoxia (> 4 hours) (Ravn et al., 2019). We investigated the behavioural, neuromuscular and CNS consequences of water immersion or exposure to N_2_ or CO_2_ gas in *Locusta migratoria* to induce anoxic coma. Previous research suggests that the gas composition in the tracheae resulting from these treatments will be markedly different. Accepting that during normal ventilation air pressures in thorax and abdomen can be locally controlled (Harrison et al., 2013), it is reasonable to assume that using N_2_ the vigorous abdominal and head pumping under respiratory distress will flush air from the tracheae inducing rapid anoxia. The action of CO_2_ to depress sensitivity to glutamate and inhibit neuromuscular transmission, e.g. in *Drosophila melanogaster* larvae (Badre et al., 2005), will rapidly immobilize the locust, stopping ventilation and delaying anoxia (measured by lactate accumulation in the hemolymph), as shown in crickets, *Acheta domesticus* (Woodring et al., 1978). Water immersion will trap air in the tracheae allowing O_2_ to be consumed until it drops below ∼2 % (∼2 kPa) (Wegener and Moratzky, 1995), triggering SD and hypometabolic paralysis. In *Schistocerca americana* under progressive hypoxia, the abdominal pumping rate of ventilation increases when the partial pressure of O_2_ (*P*_O2_) in the metathoracic ganglion drops to 5 kPa; metabolic rate decreases abruptly, indicating anoxic coma, with metathoracic *P*_O2_ below 2.5 kPa (Harrison et al., 2020). Hence, we expected that water immersion would be the most benign treatment, due to a mix of CO_2_, N_2_ and residual O_2_ in the tracheae during the anoxic coma. In addition, we expected that CO_2_ would be the most physiologically disruptive treatment, combining delayed anoxia with hypercapnia and acidosis. Given the differences in speed of coma onset and recovery with different gases (Xiao et al., 2019) and the recognition that coma is triggered by SD, an important question is whether N_2_ and CO_2_ have different effects on SD.

## Methods

### Animals

Gregarious migratory locusts, *Locusta migratoria,* were reared in a crowded colony in the animal care facility in the Biosciences Complex at Queen’s University. The colony was maintained on a 12:12 hr light-dark photoperiod with temperatures of 30 ± 1°C during light and 26 ± 1°C during dark. Locusts were fed daily with wheat grass and a dry mixture of skim milk powder, torula yeast, and bran (1:1:13 by volume). Adult locusts 3 to 6 weeks past the final moult were used for all experiments. Mass was recorded prior to any manipulation.

### Whole animal anoxia

Each anoxia treatment was performed on 10 males and 10 females. Locusts were tested in pairs, one male and one female, both taken from the same cage in the colony. For water immersion the locusts were placed in a perforated plastic container that was then submerged in a glass aquarium tank filled with de-chlorinated tap water at room temperature (∼21 °C). To monitor recovery, locusts were removed from the water and dried by removing surface water with paper towel. For gas treatment the locusts were placed in a 1 L glass filtering flask fitted with a stopper and tubing connected to a tank of either 100 % N_2_ or 100 % CO_2_. Gas was released into the flask for 1-2 minutes to replace the air before sealing the hose barb with parafilm. Locusts were removed from the flask for recovery.

Before entering a coma, locusts struggled to escape the anoxic environment by seeking an exit and attempting to jump and fly. After a variable period, activity ceased briefly before the appearance of convulsions and hindleg kicking and twitching that we took as the time of entry to coma. Locusts remained in the coma for 30 minutes before removal from the anoxic environment. We monitored recovery by noting the time it took for ventilatory movements of the abdomen to start and then for the locust to right itself, usually abruptly, and support its weight off the substrate.

### Electromyographic preparation

The hind legs and wings were removed to prevent accidental removal of the EMG electrode. The animal was restrained with plasticine and an EMG electrode (50 µm copper wire, insulated except at the tip) was placed through a pinhole in the cuticle above the spiracle on the 3^rd^ abdominal segment. The electrode was inserted ∼2 mm deep to reach an abdominal ventilatory muscle and was held in place with wax. A chlorided silver ground wire was inserted posteriorly into the thorax and was secured with a drop of wax.

Following preparation, locusts were placed into a 50 mL syringe, with one end fitted with a gas exchange and the other end open. A flow of room air was pumped through the syringe to record baseline activity for 15 minutes prior to anoxia. Then the flow was switched to either 100% CO_2_ or 100 % N_2_ gas to induce anoxia. For EMG recording with water immersion, following preparation the locust was confined in an open ventilated container to record 15 minutes of baseline activity. The locust was then placed in a 250 mL beaker half-filled with room temperature de-chlorinated water. A 200 mL beaker was placed inside the water-filled beaker to keep the animal submerged. For recovery, locusts were removed from the beaker and dried.

After the cessation of all electrical activity in the EMG trace, locusts remained in anoxia for an additional 30 minutes of coma before being returned to normoxia. Recovery was recorded for 30 minutes. We measured the time to motor pattern failure and the time to SD, indicated by the final burst of electrical activity. Recovery measures were taken as the time to excitability return and the time to motor pattern return.

### Semi-intact preparation

We used a standard preparation for investigating neural function in the thoracic ganglia (Robertson and Pearson, 1982). Locusts were pinned to a cork substrate after removing the wings, legs and pronotum. The thorax and anterior abdomen were opened with a dorsal incision and the ventral nerve cord was exposed by removing air sacs, gut, fat body, and salivary gland. The ventral diaphragm was cut away to reveal the metathoracic ganglion. This preparation was sufficient to enable recording after minimally invasive preparation for which no nerve roots were cut and the tracheal supply to the ganglia was intact. For intracellular recording, it was necessary to remove the muscles attached to the second spina between the connectives and stabilize the nervous system by supporting the meso- and metathoracic ganglia on a stainless-steel plate after cutting nerves 3, 4 and 5 on both sides of the ganglia. This minimized movement by de-efferenting the thoracic musculature. The preparation was bathed in standard locust saline (in mM: 147 NaCl, 10 KCl, 4 CaCl_2_, 3 NaOH and 10 HEPES buffer; pH = 7.2; chemicals from Sigma-Aldrich).

### Ion-sensitive electrodes

Ion-sensitive electrodes were made using silanized glass capillaries, pulled to 5-7 MΩ. For measuring [K^+^] they were filled at the tips with Potassium Ionophore I-Cocktail B (5% valinomycin; Sigma-Aldrich) and back-filled with 500 mM KCl (Rodgers et al., 2007). For [H^+^] measurement they were filled at the tips with Hydrogen ionophore I – cocktail B (10% tridodecylamine; Sigma-Aldrich) and back-filled with a solution (pH 6) of 100 mM sodium citrate and 100 mM sodium chloride (Pacey and O’Donnell, 2014). Voltage from the ion-sensitive electrode was referenced against voltage recorded with an extracellular glass microelectrode (5-7 MΩ; back-filled with 3 M KCl) positioned just adjacent. To ensure that the electrode sensitivity fell between a range of 54 to 58 mV for a 10-fold change in concentration, at room temperature, [K^+^] electrodes were calibrated using 15 mM and 150 mM KCl solutions and [H^+^] electrodes were calibrated using buffered saline at pH 6 and pH 7.

### Electrophysiology

Signals from extracellular electrodes (EMG wires or glass suction electrodes) were amplified using an AM Systems model 1700 differential AC amplifier with frequency cut-offs at 1 Hz (low) and 10 kHz (high). Signals from ion-sensitive electrodes were amplified with a Duo773 amplifier (WPI Inc., Sarasota) using a high resistance headstage for the ion-specific electrode. Intracellular recordings were made with glass microelectrodes (20-50 MΩ, back-filled with 500 mM KCl and with 3 M KCl in the electrode holder) and amplified using a model 1600 Neuroprobe amplifier (A-M Systems). All electrophysiological signals were digitized (1440A digitizer; Molecular Devices) with a sampling rate of 100 kHz and recorded using Axoscope 10.7 for later analysis using Clampfit 10.7.

### Anoxia of semi-intact preparations

Locusts were dissected inside a Plexiglas chamber (5 x 2.5 x 2 cm) that had a cork floor. For H_2_O anoxia, we completely immersed the preparation using standard locust saline, which was removed at the end of the coma with a 50 mL syringe and extension tube. For gas anoxia, the gas was supplied using a modified Pasteur pipette hooked over the end of the chamber, which could be partially sealed with electrical tape or cellophane tape (the latter enables illumination of the preparation). Positioning the tape left an aperture of approximately 1 x 1.5 cm for the electrodes to access the preparation and for the gas flow to exit. Before anoxia, an aquarium pump pumped room air through the chamber at ∼100 mL/min. To induce anoxia, flow was switched to either 100 % N_2_ or 100 % CO_2_ from pressurized tanks at ∼500 mL/min. At the end of the coma, flow was switched back to room air to flush the chamber. The duration of the coma was variable, depending on the experiment, and is noted in the Results section.

### Pharmacology

Chemicals were purchased from Sigma-Aldrich, prepared in stock solutions and frozen as aliquots for later use. Bafilomycin was purchased as 0.1 mL of a ready-made solution of 160 µM in DMSO made up to 1.6 mL of 10 µM in standard locust saline. The final concentration of DMSO has no noticeable effect in the locust nervous system (Armstrong et al., 2006). Dosage was determined from prior reports of what is effective and taking into consideration the difficulty for drugs to penetrate the blood-brain barrier and permeate the ganglion. Moreover, solutions delivered by bath application will have been diluted to approximately half the original concentration by the residual saline in the thoracic cavity. We used 10 mM ouabain to inhibit the Na^+^-K^+^-ATPase (NKA) (Van Dusen et al., 2020b) and 10 µM bafilomycin A_1_ to inhibit the V- type H^+^-ATPase (VA) (Kocmarek and O’Donnell, 2011).

### Statistical analysis

We used SigmaPlot 13 or 14 (Systat Software Inc.) to analyze the results and generate graphs. Outliers were removed prior to analysis. Depending on the experiment, sample sizes can be different because of difficulty in taking measurements from some recordings. Data were tested for normality (Shapiro-Wilk test) and equal variance (Browne-Forsythe test). Student’s t-test, ANOVA (One-Way, Two-Way and Repeated Measures as appropriate) or Kruskal-Wallis One-way ANOVA on ranks for non-parametric data were used to determine statistical significance (P < 0.05) within each measure. All-pairwise post-hoc analysis was used to determine significance between treatments within each measure: Holm-Sidak method or Bonferroni for parametric data, which are reported as mean ± standard deviation, and Tukey test or Dunn’s method for non-parametric data, which are reported as median and interquartile range (IQR). For consistency, all graphical displays of the data, whether parametric or not, are box plots showing the median and 25^th^ and 75^th^ percentiles with whiskers to the 10^th^ and 90^th^ percentiles. These are overlaid with individual data points plotted as open symbols.

## Results

### Unrestrained whole animals

Similar to crickets (Woodring et al., 1978), locusts exposed to N_2_ struggled more vigorously (i.e. more climbing, jumping and walking) than locusts exposed to CO_2_ gas (characterized by slower movements, in some instances no movement at all). Immersed locusts struggled the most intensely and for a longer period. Considering the full dataset of 60 locusts, none of the measures was affected by sex (Kruskal-Wallis ANOVA: time to succumb P = 0.89; time to ventilate P = 0.51; time to stand P = 0.77). Hence, except as noted below, we combined the male and female data in the following analyses.

The time to enter a coma (**Fig. 1A**) depended on the method of anoxia (Kruskal-Wallis ANOVA: P < 0.001; n = 19 H_2_O, 20 N_2_, 20 CO_2_). Immersion in water took the longest time at 2.1 (1.9-2.5) mins, followed by nitrogen at 0.9 (0.6-1.0) mins and then carbon dioxide at 0.3 (0.3-0.4) mins (Dunn’s: H_2_O vs CO_2_ – P < 0.001; H_2_O vs N_2_ – P < 0.001; N_2_ vs CO_2_ – P = 0.007).

**Figure 1.**
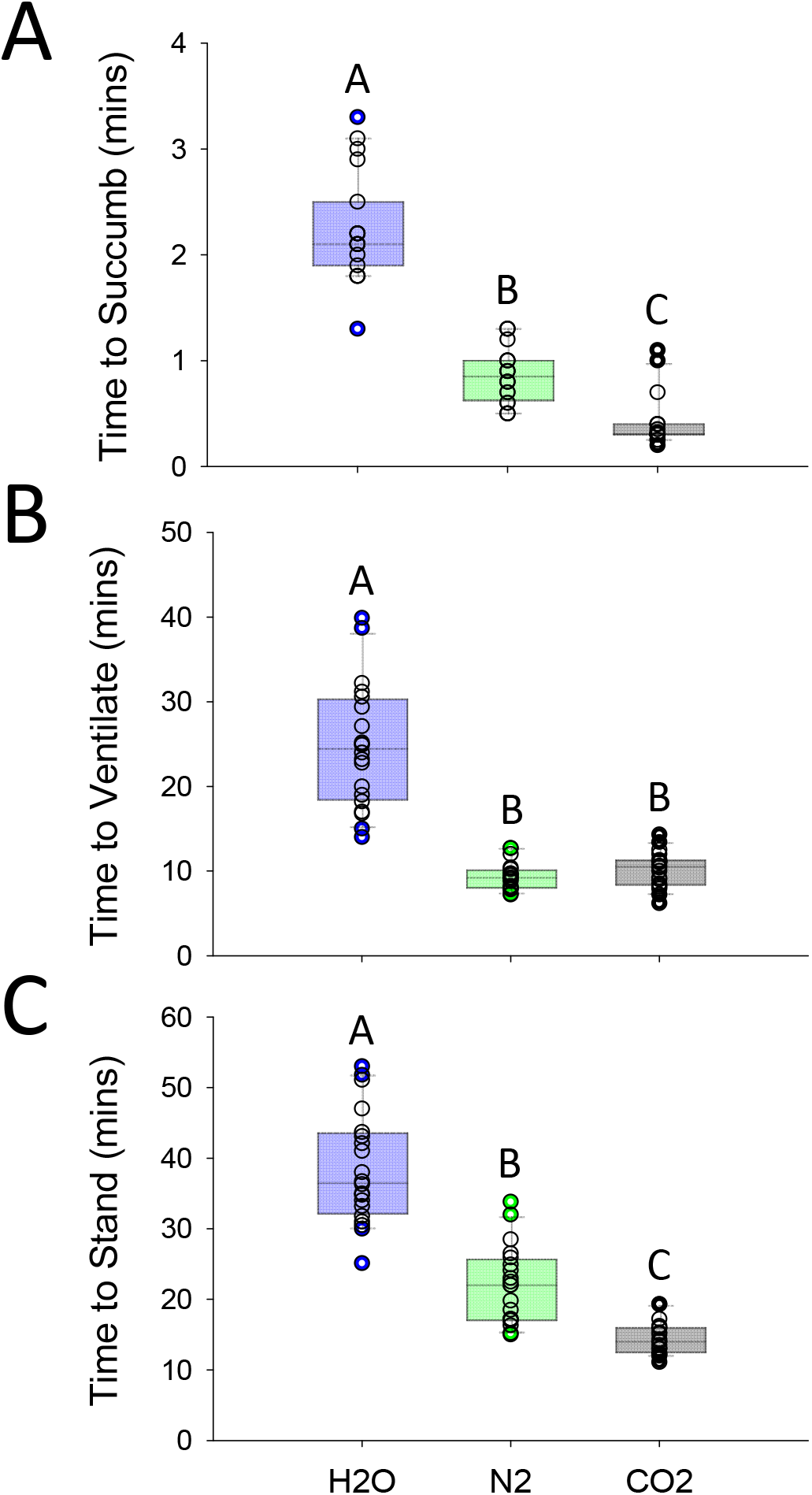
Timing of entry to, and recovery from, anoxic coma of whole animals depends on method of anoxia. **A.** Time taken for locusts, including males and females, to succumb to anoxia and enter a coma after immersion in water or exposure to nitrogen or carbon dioxide gas. **B.** Time taken for ventilation movements of the abdomen to start after return to air. **C.** Time taken for the locust to stand after return to air. Box plots indicate the median, 25^th^ and 75^th^ percentiles with whiskers to the 10^th^ and 90^th^ percentiles. Individual data points plotted as open symbols. Statistically significant differences indicated with different letters.

The time to recover ventilation (**Fig. 1B**) depended on the method of anoxia (Kruskal-Wallis ANOVA: P < 0.001; n = 20, 20, 20). Immersion in water took the longest time at 24.5 (18.4-30.3) mins, but there was no statistical difference between nitrogen at 9.2 (8.0-10.1) mins and carbon dioxide at 10.5 (8.4-11.3) mins (Dunn’s: H_2_O vs CO_2_ – P < 0.001; H_2_O vs N_2_ – P < 0.001; N_2_ vs CO_2_ – P = 0.61).

The time to stand (**Fig. 1C**) depended on the method of anoxia (Kruskal-Wallis ANOVA: P < 0.001; n = 20, 20, 20). Immersion in water took the longest time at 36.5 (32.2-43.6) mins, followed by nitrogen at 22.0 (17.1-25.7) mins and carbon dioxide at 14.0 (8.4-11.3) mins (Dunn’s: H_2_O vs CO_2_ – P < 0.001; H_2_O vs N_2_ – P < 0.001; N_2_ vs CO_2_ – P = 0.004).

The above statistical treatment of the data was hampered by the lack of normality and/or equal variance in the full dataset. Given that sex differences in recovery from anoxia have been reported (e.g., for *Chortoicetes terminifera*, Robertson et al. 2019) we looked for any effects of sex within measures separately. We found that males took longer to recover ventilation after nitrogen anoxia (male 10.4 ± 1.6 mins; female 8.3 ± 0.8 mins; Student’s t-test P = 0.001; n = 10, 10) and females took longer to stand after carbon dioxide anoxia (male 13.2 ± 1.5 mins; female 15.7 ± 2.3 mins; Student’s t-test P = 0.009; n = 10, 10). There were no other differences.

#### Summary

Intact, unrestrained locusts entered a coma and recovered the ability to stand most rapidly under CO_2_ anoxia. To obtain more objective measures of the neuromuscular basis for this difference we recorded ventilatory muscle activity with anoxia in intact, restrained locusts.

### Electromyographic recording

To characterize neuromuscular failure and recovery due to anoxia, we collected a dataset from 30 locusts, using males only to reduce variability. We monitored coma induction at two well-defined time points: the cessation of ventilatory muscle motor patterning and the end of the final burst of unpatterned activity before neuromuscular silence indicating entry to coma (**Fig. 2A**). The method of anoxia influenced the time to both measures: time to motor pattern failure (Kruskal-Wallis ANOVA: P < 0.001, n = 10, 10, 10) and time to coma (Kruskal-Wallis ANOVA: P < 0.001, n = 10 per treatment). Both measures show that the CO_2_ treatment significantly decreased induction times. The time to motor pattern failure was shorter with CO_2_ treatments compared to N_2_ (Tukey: P = 0.003), and compared to H_2_O treatments (Tukey Test, P < 0.001) (**Fig. 2B**) (H_2_O 2.7 (1.5-8.5) mins; N_2_ 1.7 (1.5-2.2) mins; CO_2_ 0.3 (0.2-0.3) mins). Similarly, coma occurred sooner with CO_2_ treatments compared to N_2_ treatments (Tukey: P = 0.002), and H_2_O treatments (Tukey: P < 0.001) (**Fig. 2C**) (H_2_O 4.9 (2.6-10.0) mins; N_2_ 3.6 (3.0-4.4); CO_2_ 0.7 (0.6-1.0) mins).

**Figure 2.**
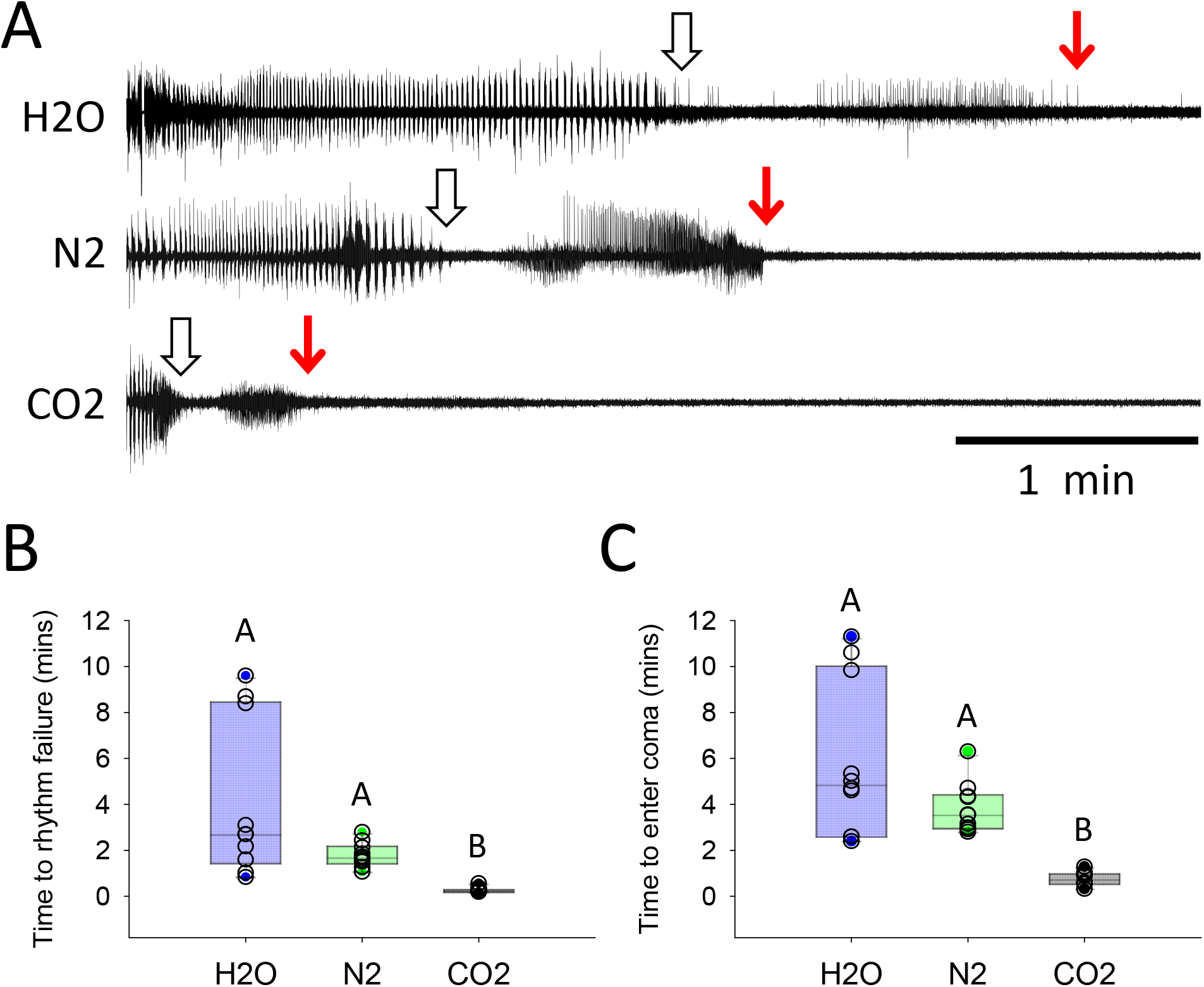
euromuscular failure is more rapid with CO_2_ anoxia. **A.** Sample EMG traces from male locusts during exposure to different agents of anoxia starting at the beginning of the recording. Open arrows indicate failure of rhythmic ventilatory activity; red arrows indicate the beginning of electrical silence characteristic of coma. **B.** Time to rhythm failure is shorter with CO_2_. **C.** Time to enter coma is shorter with CO_2_. Box plots indicate the median, 25^th^ and 75^th^ percentiles with whiskers to the 10^th^ and 90^th^ percentiles. Individual data points plotted as open symbols. Statistically significant differences indicated with different letters.

Neuromuscular recovery was characterized by measuring the time to the return of electrical excitability and the time to return of motor patterning (**Fig. 3A**). In a few preparations, there was some difficulty, and thus a subjective element, in determining precisely when tonic activity changed to consistent patterned activity because this was not a clear-cut transition. However, this inaccuracy is contained within the large variances due simply to individual variation. Recovery of excitability depended on the method of anoxia (One-Way ANOVA: P < 0.001; n = 10, 10, 10). H_2_O delayed recovery compared to N_2_ (Holm-Sidak: P < 0.001) and compared to CO_2_ treatments (Holm-Sidak: P < 0.001). CO_2_ delayed recovery compared to N_2_ (Holm-Sidak: P = 0.037) (**Fig. 3B**) (H_2_O 12.8 ± 3.5 mins; N_2_ 4.7 ± 1.9 mins; CO_2_ 7.3 ± 2.5 mins). Recovery of motor patterning also depended on the method of anoxia (One-Way ANOVA: P < 0.001; n = 10, 10, 10) and was delayed with H_2_O compared to N_2_ (Holm-Sidak: P<0.001) and CO_2_ (Holm-Sidak: P < 0.001) (**Fig. 3C**) (H_2_O 16.8 ± 3.6 mins; N_2_ 7.6 ± 3.1 mins; CO_2_ 10.3 ± 3.5 mins). The motor pattern changed after recovery. Prior to anoxia the duration of motor bursts was 0.44 ± 0.24 s with a frequency of 1.03 ± 0.29 Hz. After recovery the burst duration increased though the difference in fold-increase due to the method of anoxia was only marginally significant (One-Way ANOVA: P = 0.05; H_2_O 1.4 ± 0.8-fold; N_2_ 2.2 ± 1.0-fold; CO_2_ 1.2 ± 0.7-fold) (**Fig. 3E**). The method of anoxia did influence the fold-change of the pattern frequency, which was reduced by gas anoxia compared to water immersion (One-Way ANOVA on transformed data (logx): P < 0.001; H_2_O 1.03 ± 0.35-fold; N_2_ 0.34 ±0.15 fold; CO_2_ 0.30 ± 0.11-fold) (**Fig. 3F**).

**Figure 3.**
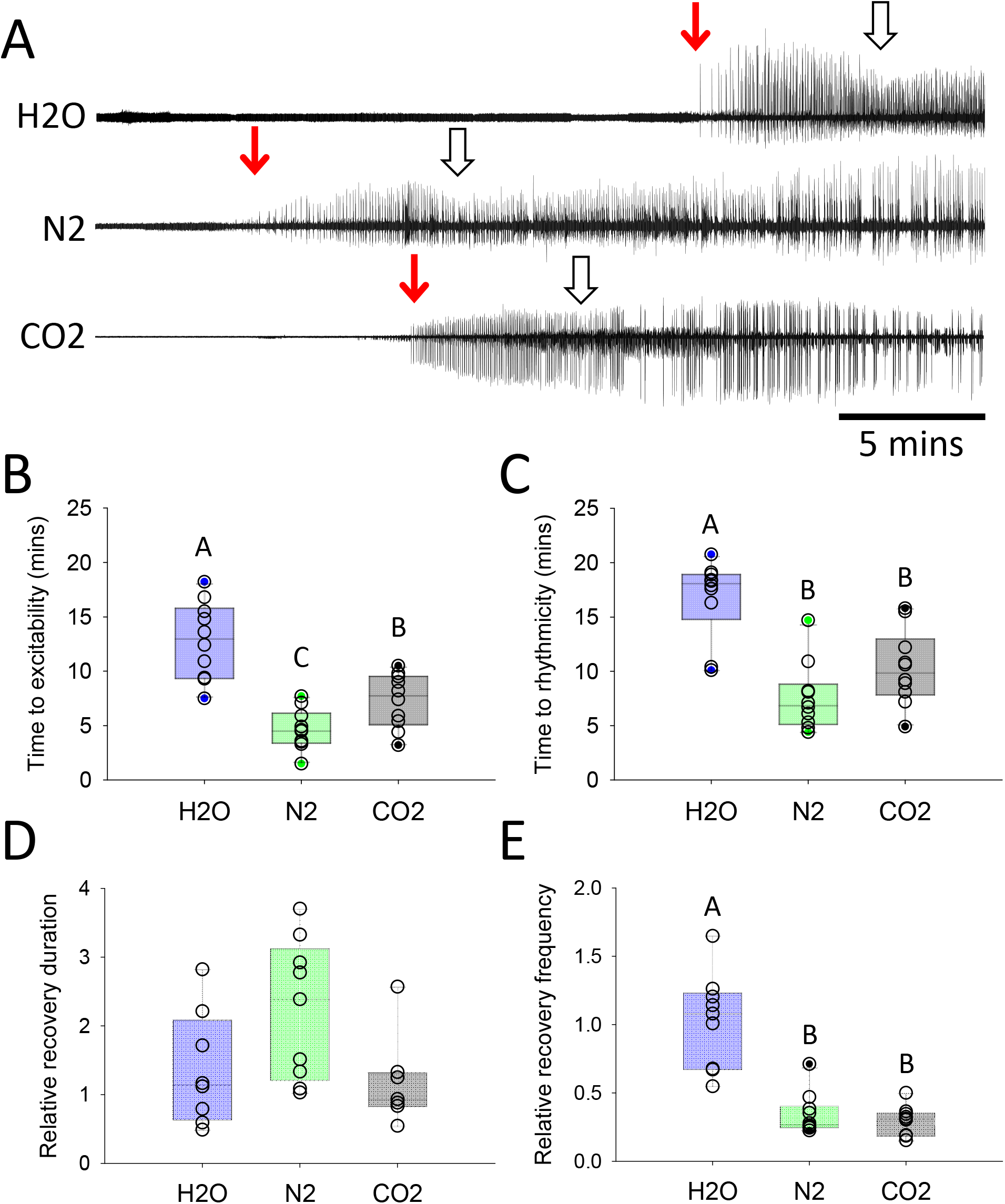
Neuromuscular recovery depends on method of anoxia. **A.** Sample EMG traces from male locusts recovering after exposure to different agents of anoxia. Air returns at the beginning of the recording. Red arrows indicate the recovery of electrical excitability; open arrows indicate recovery of rhythmic ventilatory activity. Note that the recovery of ventilatory motor patterning can be difficult to identify, particularly in these compressed traces. **B.** Time to recover excitability. **C.** Time to recover rhythmicity. **D.** Relative duration of ventilatory motor bursts compared with starting values. **E.** Relative frequency of the ventilatory motor pattern compared with starting values. Box plots indicate the median, 25^th^ and 75^th^ percentiles with whiskers to the 10^th^ and 90^th^ percentiles. Individual data points plotted as open symbols. Statistically significant differences indicated with different letters.

#### Summary

Neuromuscular measures of entry to coma were qualitatively like the behavioural measures. However, in contrast with the behavioural measures, we found that recovery of neural function measured electromyographically, which neglects any contribution of muscle contractility and strength, was most rapid under N_2_ anoxia. To investigate the characteristics of SD, we recorded the transperineurial potential of the metathoracic ganglion during anoxia of semi-intact preparations.

### Transperineurial potential recording

SD is characterized by an abrupt negative shift in the extracellular DC potential. In locusts this is recorded from the interstitial space of the ganglion relative to the bathing saline and is equivalent to the transperineurial potential (TPP) across the blood brain barrier (BBB) (Robertson et al., 2020; Schofield and Treherne, 1984). To characterize the dynamics of SD we recorded the TPP simultaneously with a nerve recording of ventilatory activity and an EMG recording from muscles controlling the hindwing (**Fig. 4**). In 10 locusts for each method of anoxia we measured the time for the ventilatory rhythm to fail (**Fig. 5A**) and the time to the onset of SD taken at the half-amplitude of the negative DC shift (**Fig. 5B**). After ∼1 min in the coma (timed from SD onset) the preparation was returned to normoxia and we measured the time for the return of ventilatory motor patterning (**Fig. 5C**) and the time for the TPP to return to normal taken at the half-amplitude of the TPP recovery (**Fig. 5D**). To characterize the negative DC shift we measured its amplitude (**Fig. 5E**) and the slope of the TPP recovery (**Fig. 5F**).

**Figure 4.**
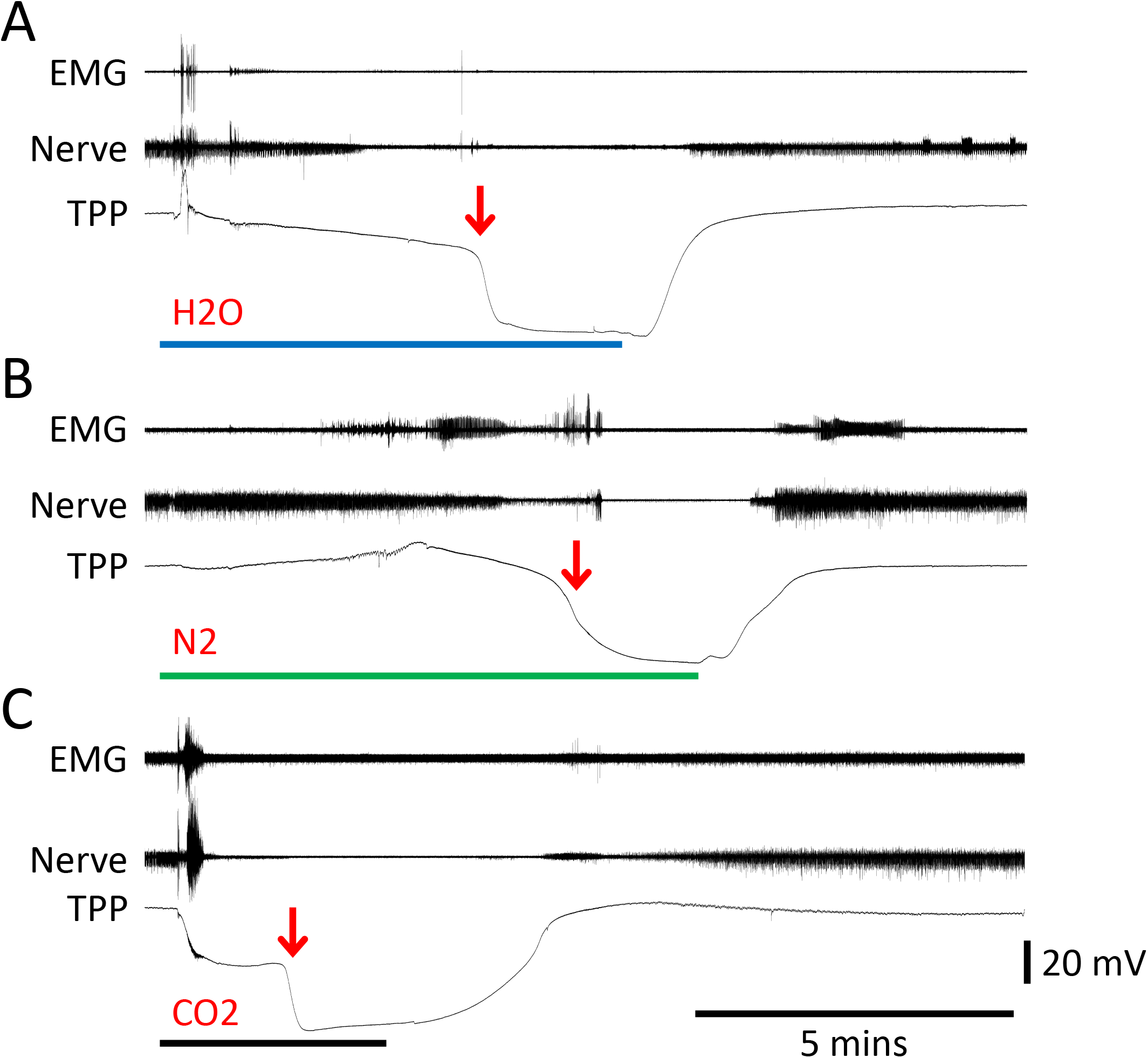
CNS shutdown and recovery are associated with characteristic changes of the transperineurial potential (TPP). Recordings made during coma onset and recovery due to treatment with **A.** H_2_O, **B.** N_2_ and **C.** CO_2_. Treatment duration is indicated by the lines under the traces. SD onset is indicated by the red arrow. The voltage scale bar is for the TPP trace and is the same for all panels.

**Figure 5.**
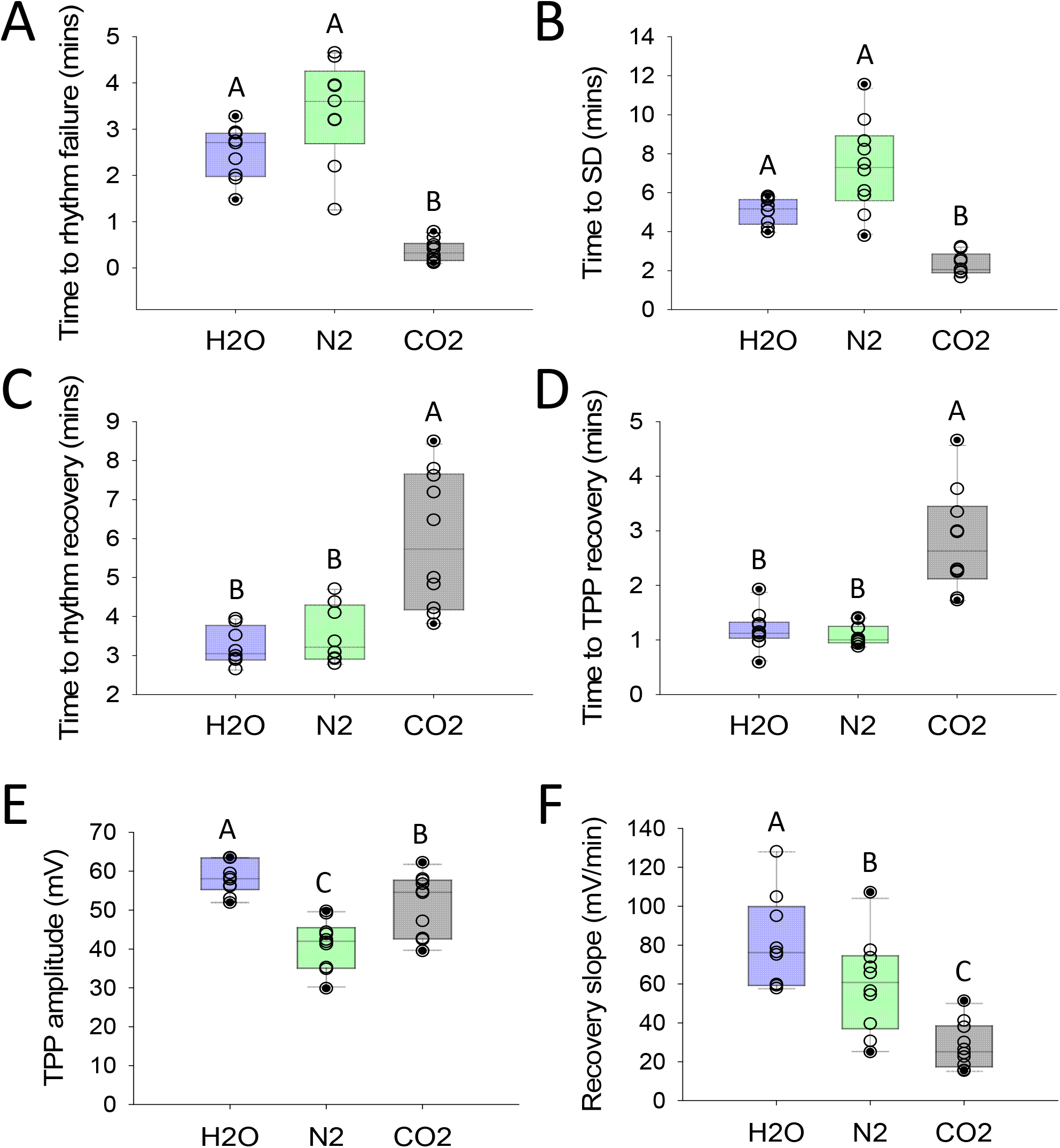
CNS shutdown is faster, and recovery is slower with CO_2_ anoxia. Entry to and recovery from a ∼1 min coma induced different methods of anoxia characterized by **A.** Time to rhythm failure, **B.**Time to SD (negative DC shift), **C.** Time to rhythm recovery, **D.** Time to TPP recovery (postitive DC shift), **E.** Amplitude of the TPP shift, **F.** Slope of the TPP returning to normal. Box plots indicate the median, 25^th^ and 75^th^ percentiles with whiskers to the 10^th^ and 90^th^ percentiles. Individual data points plotted as open symbols. Statistically significant differences indicated with different letters.

The method of anoxia affected the time for motor patterning to fail (Kruskal-Wallis ANOVA: P < 0.001; n = 10 H_2_O, 9 N_2_, 10 CO_2_). Time to rhythm failure was shorter with CO_2_ than with either H_2_O (Dunn’s: P = 0.005) or N_2_ (Dunn’s: P < 0.001) but there was no difference between H_2_O and N_2_ (Dunn’s: P = 0.48). (H_2_O 2.7 (2.0-2.9) mins; N_2_ 3.6 (2.7-4.3) mins; CO_2_ 0.3 (0.2-0.5) mins). Similarly, the method of anoxia affected the time to enter a coma (Kruskal-Wallis ANOVA: P < 0.001; n = 10 H_2_O, 10 N_2_, 9 CO_2_). Time to SD was shorter with CO_2_ than with either H_2_O (Dunn’s: P = 0.01) or N_2_ (Dunn’s: P < 0.001) but there was no difference between H_2_O and N_2_ (Dunn’s: P = 0.25). (H_2_O 5.2 (4.4-5.7) mins; N_2_ 7.3 (5.6-8.9) mins; CO_2_ 2.1 (1.9-2.9) mins).

The method of anoxia affected the time for motor patterning to recover (Kruskal-Wallis ANOVA: P < 0.001; n = 8 H_2_O, 8 N_2_, 10 CO_2_). Time to rhythm recovery was longer with CO_2_ than with either H_2_O (Dunn’s: P = 0.002) or N_2_ (Dunn’s: P = 0.02) but there was no difference between H_2_O and N_2_ (Dunn’s: P = 1.0). (H_2_O 3.1 (2.9-3.8) mins; N_2_ 3.2 (2.9-4.3) mins; CO_2_ 5.7 (4.2-7.7) mins). Similarly, the method of anoxia affected the time for the TPP to recover (Kruskal-Wallis ANOVA: P < 0.001; n = 10, 10, 10). Time to TPP recovery was longer with CO_2_ than with either H_2_O (Tukey: P = 0.002) or N_2_ (Tukey: P < 0.01) but there was no difference between H_2_O and N_2_ (Tukey: P = 1.0). (H_2_O 1.1 (1.0-1.3) mins; N_2_ 1.0 (1.0-1.3) mins; CO_2_ 2.6 (2.1-3.5) mins).

The amplitude of the negative DC shift was dependent on the method of anoxia (One Way ANOVA: P < 0.001; n = 10, 10, 10). TPP amplitude with H_2_O was larger than with N_2_ (Holm-Sidak: P < 0.001) or with CO_2_ (Holm-Sidak: P = 0.02). There was also a difference in TPP amplitude between N_2_ and CO_2_ (Holm-Sidak: P = 0.002). (H_2_O 58 ± 4.2 mV; N_2_ 41 ± 6.3 mV; CO_2_ 52 ± 7.9 mV). The method of anoxia also affected the slope of the TPP recovery trajectory (One Way ANOVA: P < 0.001; n = 10, 10, 10). This slope was larger with H_2_O than with N_2_ (Holm-Sidak: P = 0.03) or with CO_2_ (Holm-Sidak: P < 0.001). There was also a difference in the TPP recovery slope between N_2_ and CO_2_ (Holm-Sidak: P = 0.004). (H_2_O 82 ± 24 mV/min; N_2_ 60 ± 24 mV/min; CO_2_ 28 ± 12 mV/min).

#### Summary

CO_2_ had a strong effect on the timing of neural failure and entry to anoxic coma (earlier) and recovery on return to normoxia (later) but there was no difference between H_2_O and N_2_. However, the size and shape of the negative DC shift of the TPP was different in each of the treatments. Delivery of 100 % CO_2_ gas is expected to generate a rapid hypercapnic acidosis as the gas is delivered directly to the CNS through the tracheoles while the locust continues to ventilate. To confirm this, in the next experiments we measured ion concentrations in the interstitial space of the metathoracic ganglion.

### Measurement of extracellular ion concentrations

Given that there was no difference between H_2_O and N_2_ in the timing of SD and recovery we compared extracellular ion concentration changes only for N_2_ and CO_2_ anoxia. We measured interstitial pH (pH_o_) in 6 male locusts for each gas. The onset of N_2_ provoked an immediate increase in the frequency of the ventilatory rhythm but had no effect on pH_o_ (**Fig. 6A**). In contrast, the onset of CO_2_ caused an abrupt decrease in pH_o_ and the increase in the ventilatory rhythm frequency was transient before a large burst of unpatterned nerve activity leading to electrical silence (**Fig. 6B**). Nevertheless, pH_o_ decreased for both gases during the coma (**Fig. 6C,D**). After the return to normoxia, pH_o_ recovery took longer than the recovery of the TPP. In addition, there was a transient decrease in pH_o_ around the time that neuronal excitability recovered (**Fig. 6C,D**). At the start of recovery, pH_o_ increased by 0.26 ± 0.14 units per minute but transiently decreased by −0.20 ± 0.18 units per minute when nerve activity resumed (**Fig. 6E**). Both the nature of the gas and the time of measurement (before and after SD) had effects on pH_o_ (**Fig. 6F**) and there was a significant interaction (Two-way ANOVA: P_gas_ = 0.028; P_time_ < 0.001; P_gas x time_ = 0.01). pH_o_ before anoxia was 7.3 ± 0.4 (N_2_) and 7.4 ± 0.3 (CO_2_) (Holm-Sidak: P = 0.75) and reached a peak during the coma of 6.8 ± 0.4 for N_2_, and significantly lower 6.0 ± 0.3 for CO_2_ (Holm-Sidak: P = 0.001). The pH_o_ reduction was significant for both gases (Holm-Sidak: P = 0.02 for N_2_; P < 0.001 for CO_2_).

**Figure 6.**
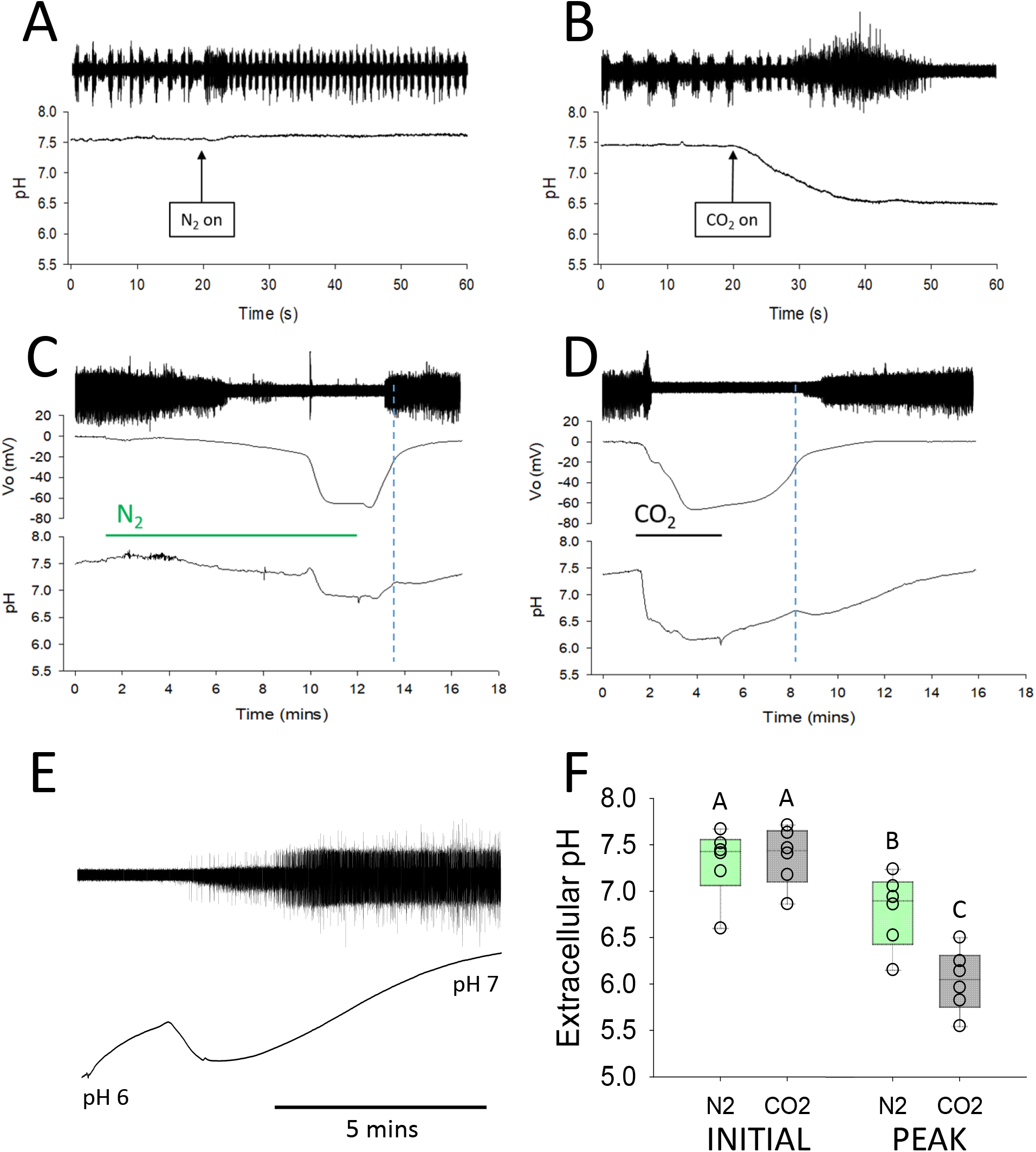
CO_2_ onset induces abrupt interstitial pH decrease. **A.** Recording ventilatory rhythm from a median nerve (top) and interstitial pH_o_ from the neuropil (bottom) at the onset of N_2_ anoxia. **B.** Same for the onset of CO_2_ anoxia. **C.** Nerve recording, negative DC shift (V_o_) and pH_o_ before, during and after N_2_ anoxia with a 1 minute coma. **D.** Same for CO_2_ anoxia. Timing of gas delivery indicated by lines under the traces. Note the discontinuity of pH_o_ recovery indicated by the dotted lines. **E.** In a different preparation, discontinuous pH_o_ recovery is coincident with recovery of excitability. The traces start 30 s after the return of air following a 1 min CO_2_ anoxia. **F.** Comparison of pH changes with N_2_ and CO_2_ anoxia. Box plots indicate the median, 25^th^ and 75^th^ percentiles with whiskers to the 10^th^ and 90^th^ percentiles. Individual data points plotted as open symbols. Statistically significant differences indicated with different letters.

We recorded [K^+^]_o_ in 7 male locusts for each gas and quantified it at three time points: prior to gas onset (Initial [K^+^]_o_); at the approximate start of the abrupt increase (Ignition [K^+^]_o_); and during the coma (Plateau [K^+^]_o_). Similar to the pH experiments, the onset of N_2_ provoked an immediate increase in the frequency of the ventilatory rhythm but had no effect on [K^+^]_o_ (**Fig. 7A**). In contrast, the onset of CO_2_ caused an abrupt increase in [K^+^]_o_ and the increase in the ventilatory rhythm frequency was transient before a large burst of unpatterned nerve activity leading to electrical silence (**Fig. 7B**). SD was associated with the characteristic surge of [K^+^]_o_ for both gases (**Fig. 7C,D**). The [K^+^]_o_ values during the surge were not affected by the nature of the gas (**Fig. 7E,F,G**). Initial [K^+^]_o_ was 13.1 ± 2.6 mM for N_2_ and 13.4 ± 2.3 mM for CO_2_ (Student’s t test: P = 0.77); Ignition [K^+^]_o_ was 31.5 ± 5.8 mM for N_2_ and 28.7 ± 6.6 mM for CO2 (Student’s t test: P = 0.42); Plateau [K^+^]_o_ it was 99.0 ± 22.0 mM for N_2_ and 97.9 ± 11.8 for CO_2_ (Student’s t test: P = 0.91).

**Figure 7.**
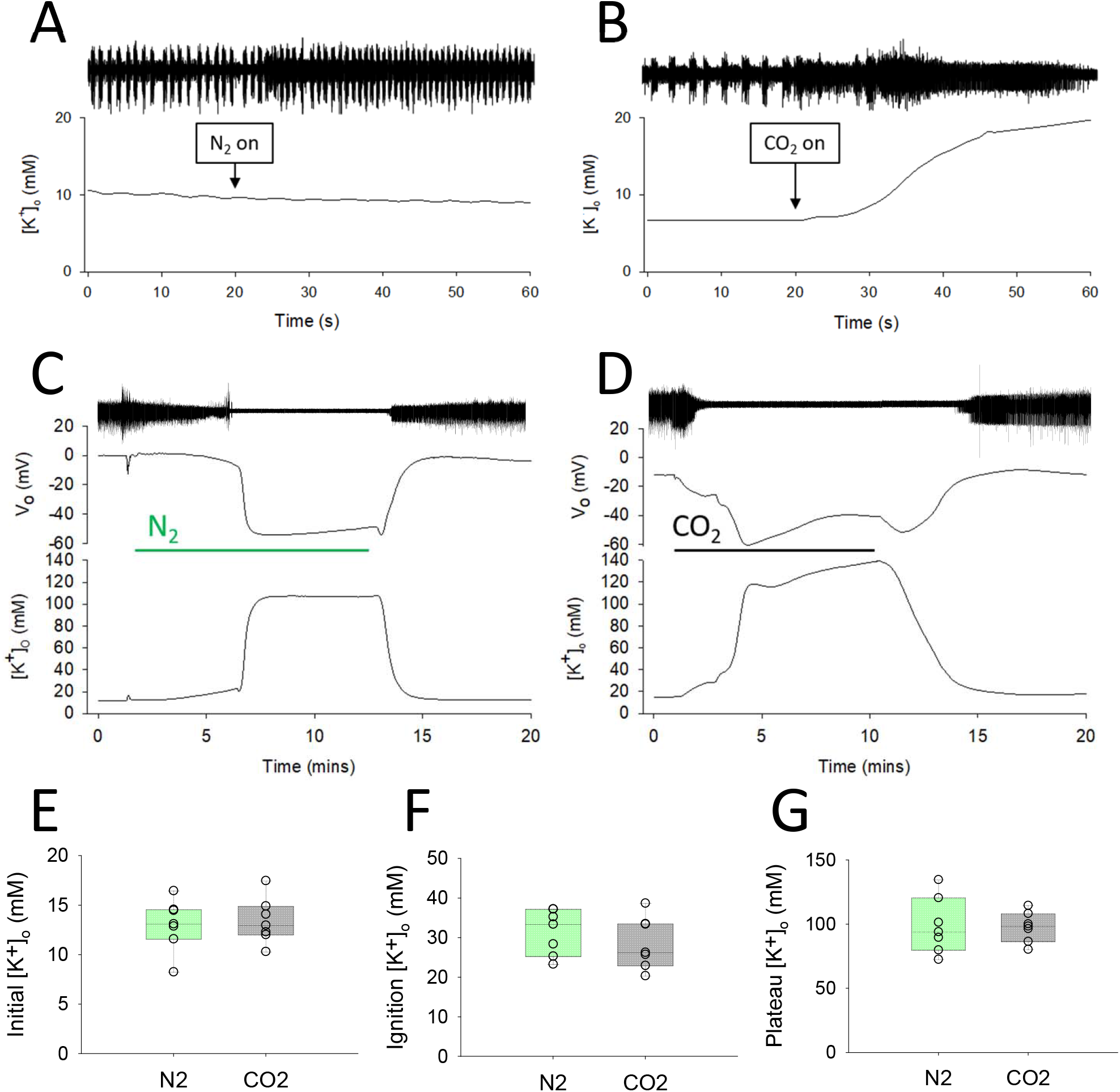
CO_2_ onset induces abrupt [K^+^]_o_ increase. **A.** Recording ventilatory rhythm from a median nerve (top) and [K^+^]_o_ from the neuropil (bottom) at the onset of N_2_ anoxia. **B.** Same for the onset of CO_2_ anoxia. **C.** Nerve recording, negative DC shift (V_o_) and [K^+^]_o_ before, during and after N_2_ anoxia with a 1 minute coma. **D.** Same for CO_2_ anoxia. Timing of gas delivery indicated by lines under the traces. **E, F, G.** Comparison of [K^+^]_o_ changes with N_2_ and CO_2_ anoxia. Box plots indicate the median, 25^th^ and 75^th^ percentiles with whiskers to the 10^th^ and 90^th^ percentiles. Individual data points plotted as open symbols.

#### Summary

Delivery of CO_2_ had immediate effects to decrease pH_o_ and increase [K^+^]_o_, which were not evident with N_2_. The surge of [K^+^]_o_ as a consequence of SD was not quantitatively affected by the nature of the gas, though the overall shape could vary. The negative DC shift associated with anoxia-induced SD propagates throughout the neuropil and on return to air there is a variable postanoxic negativity (PAN), which has been attributed to NKA (e.g. (Spong et al., 2016b)). To determine if the SD propagation rate and recovery was affected by the gas, we investigated these features in the next experiments.

### Propagation and Postanoxic Negativity

To investigate SD propagation and the PAN, we recorded TPP at two locations in 14 male locusts each of which had a N_2_ and a CO_2_ coma (5 mins duration after SD onset) with presentation order alternating between animals. In addition, to determine the consequences of the more invasive preparation required for intracellular recording, 6 of the 14 were minimally dissected and 8 were prepared for intracellular recording (i.e. with nerve roots cut and the meso- and metathoracic ganglia supported on a metal plate).

For both N_2_ and CO_2_ it was possible to measure a latency between the negative DC shifts recorded at different locations while the immediate effects of gas onset and return of air were simultaneous (**Fig 8A,B**). This latency was highly variable because it depends on the relative positions of the electrodes and the location of SD ignition, which could not be controlled. The latency was 23.5 ± 14.8 s (n = 28) with no effect of the method of anoxia or of cutting nerve roots and no interaction (Two-way ANOVA: P_gas_ = 0.22; P_cut_ = 0.42; P_gas x cut_ = 0.22). Electrode separation was ∼0.75 mm, giving a propagation speed of ∼2 mm/min. On the other hand, cutting the nerve roots and the gas did have an effect on the PAN amplitude (Two-way RM ANOVA: P_gas_ = 0.004; P_cut_, 0.002; P_gas x cut_ = 0.03). Cutting had a greater effect with CO_2_, dropping PAN amplitude from 6.2 ± 0.74 mV to 1.5 ± 0.64 mV (Bonferroni: P < 0.001) whereas with N_2_ cutting dropped PAN amplitude from 3.8 ± 0.74 mV to 1.2 ± 0.64 mV (Bonferroni: P = 0.016). In minimally dissected preparations there was a significant effect of the gas (Bonferroni: P = 0.002) but not in preparations with the nerve roots cut (Bonferroni: P = 0.49).

**Figure 8.**
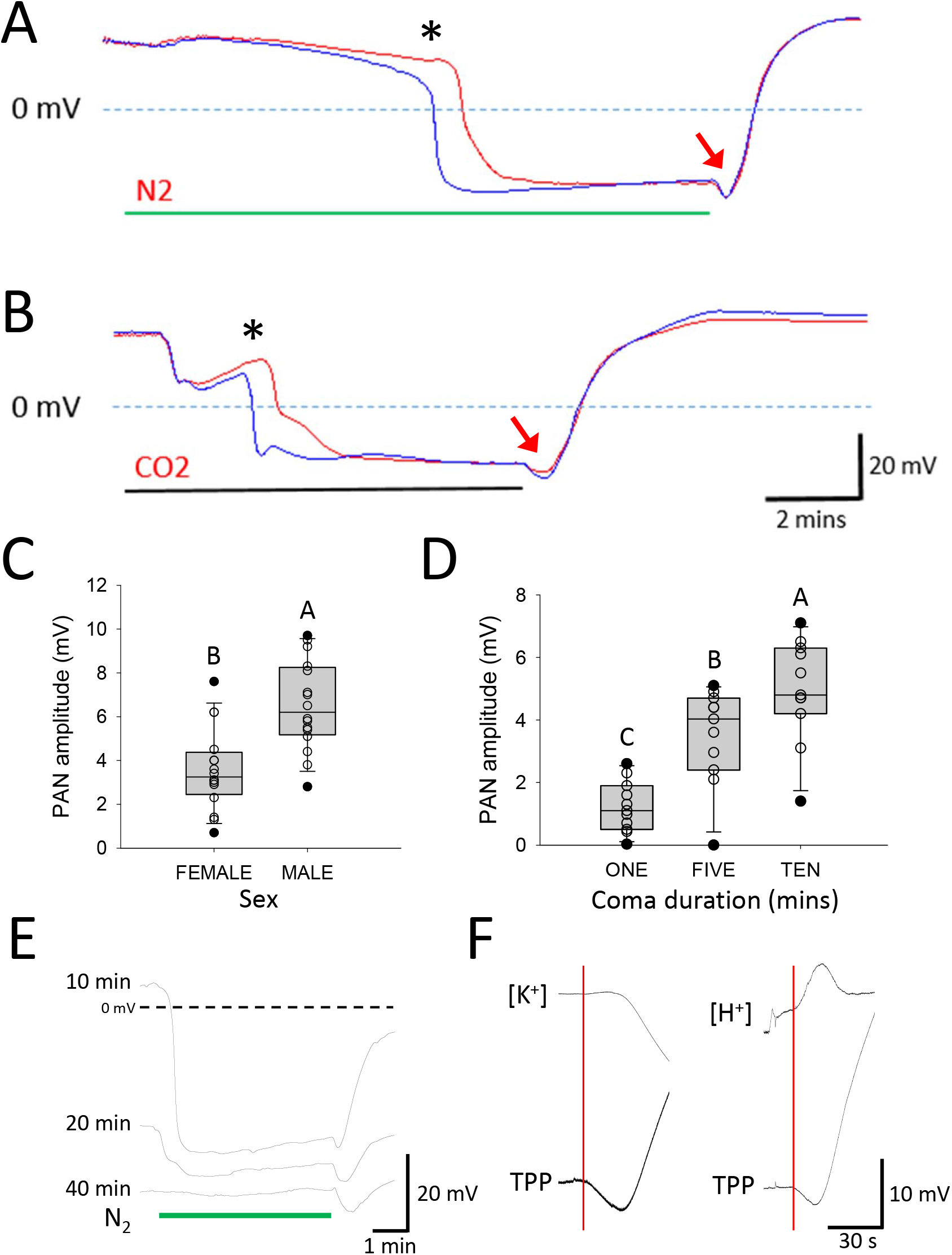
SD propagation and the postanoxic negativity. **A.** Overlaid TPP recordings taken from two locations separated by ∼0.75 mm during a 5 min N_2_ anoxia. The asterisk indicates the onset of SD, which is not simultaneous at the two locations suggesting propagation between the electrodes. The red arrow indicates a prominent postanoxic negativity (PAN) when air returned. **B.** Same for CO_2_ anoxia. Note in **A** and **B** that the effects of gas on and gas off are simultaneous at the two recording sites. **C.** PAN was larger in males. **D.** PAN was larger with longer durations of coma. **E.** TPP recordings during 5 mins of N_2_ coma at different times (10, 20 and 40 mins) after bathing the preparation with 10 mM ouabain. The dotted line at 0 mV relates only to the 10 min trace. The two other traces have each been negatively displaced by ∼5 mV for clarity but are at the same scale. Note that after 40 mins of ouabain there is no negative shift of TPP with N_2_ onset but the PAN on return to air remains and is larger than at previous time points. **F.** Sample voltage traces from the ion-sensitive electrodes and TPP showing that the PAN is not associated with any change in [K^+^]_o_ but is associated with a transient increase of [H^+^]_o_. The red vertical lines are aligned with the onset of the PAN. Box plots in C and D indicate the median, 25^th^ and 75^th^ percentiles with whiskers to the 10^th^ and 90^th^ percentiles. Individual data points plotted as open symbols. Statistically significant differences indicated with different letters.

In a separate project investigating the effects of saline additives (glucose and trehalose; manuscript in preparation), we measured PAN amplitude in 36 minimally dissected locusts (18 male and 18 female) after 10 mins of N_2_ coma. There was no effect of the saline additives but a strong effect of sex with no interaction (Two-way ANOVA: P_sex_ < 0.001; P_saline_ = 0.84; P_sex x saline_ = 0.54). PAN amplitude was 7.2 ± 0.58 mV in males and 3.5 ± 0.62 in females (Holm-Sidak: P < 0.001) (**Fig. 8C**).

The length of time in a coma is known to affect the recovery from N_2_-induced SD (Van Dusen et al., 2020b). To investigate the effect of coma duration on PAN amplitude, 12 locusts were each given N_2_ comas with durations of 1, 5 and 10 mins from the onset of SD, with presentation order arranged so that all combinations of coma duration and presentation order were represented. There was a strong effect of coma duration on PAN amplitude but no effect of presentation order (Two-way ANOVA: P_duration_ < 0.001; P_order_ = 0.31; P_duration x order_ = 0.49). The largest PAN amplitude occurred after 10 mins of coma (5.4 ± 2.3 mV), then 5 mins (4.0 ± 2.2) and 1 min (1.6 ± 1.5 mV) (Holm-Sidak: TEN vs. ONE P < 0.001; FIVE vs. ONE P = 0.001; TEN vs. FIVE P = 0.03) (**Fig. 8D**).

It has been suggested that the PAN is associated with re-activation of NKA when normoxic mitochondrial operation resumes (Spong et al., 2016b). To test this, we investigated the effect of the NKA inhibitor ouabain on the PAN in 6 male locusts given repeated 5 min N_2_ comas. Prolonged exposure to 10 mM ouabain in a minimally dissected preparation results in a gradual negative shift of the TPP from ∼ 15 mV to ∼ −40 mV, which is taken to indicate the transition from 100 % NKA function to 0 % NKA function (Van Dusen et al., 2020b). In none of the locusts was the PAN eradicated or even reduced; indeed, it increased in amplitude and duration (**Fig. 8E**). To gain insight into the underlying mechanism, we re-examined the recordings made with ion-sensitive electrodes (see above) and found that in none of the [K^+^]_o_ recordings was there any indication of [K^+^]_o_ changes coincident with the PAN. However in 8 of the 12 preparations for [H^+^]_o_ recording we could detect an increase in [H^+^]_o_ equivalent to a mean pH_o_ decrease of 0.04 ± 0.03 units (**Fig. 8F**).

#### Summary

The nature of the gas used for anoxia had no effect on SD propagation but in minimally dissected preparations the PAN was larger after CO_2_ anoxia than after N_2_ anoxia. The PAN was also larger in males and positively correlated with coma duration. We could not link the PAN to NKA function, but it was associated with a drop in pH_o_. One possibility is that the PAN represents the electrogenic effect of a proton pump (VA). We next tested that using the VA inhibitor bafilomycin.

### Bafilomycin

To investigate the effect on the PAN of inhibiting VA we recorded TPP during repeated 10-minute duration N_2_ comas. For 9 control locusts there was no intervention and for 10 experimental locusts 200 µL of 10 µM bafilomycin was added to the bathing saline after the first coma. Bafilomycin had two striking effects. First, it generated a positive DC shift in the TPP of 8.6 ± 2.9 mV, which developed over 9.3 ± 1.4 mins (n = 10). Second, it eradicated the PAN at the return to normoxia (**Fig. 9A**; example from a CO_2_ coma). This is clear after ouabain pre-treatment, which accentuated the PAN; subsequent bafilomycin eradicated it (**Fig. 9B**; example from a N_2_ coma). The PAN amplitude was reduced by bafilomycin but not by repeated anoxia (Two-way RM ANOVA: P_drug_ < 0.001; P_anoxia_ = 0.007; P_drug x anoxia_ = 0.011). In control preparations the first PAN amplitude was 4.9 ± 1.3 mV and the second was 4.8 ± 1.3 mV (Bonferroni: P = 0.88), whereas in experimental preparations the first PAN amplitude was 3.9 ± 2.1 mV and the second, after bafilomycin, was 1.3 ± 1.2 mV (Bonferroni: P < 0.001). There was no difference between PANs of first anoxias (Bonferroni: P = 0.15) but there was between PANs of second anoxias (Bonferroni: P < 0.001) (**Fig. 9C**).

**Figure 9.**
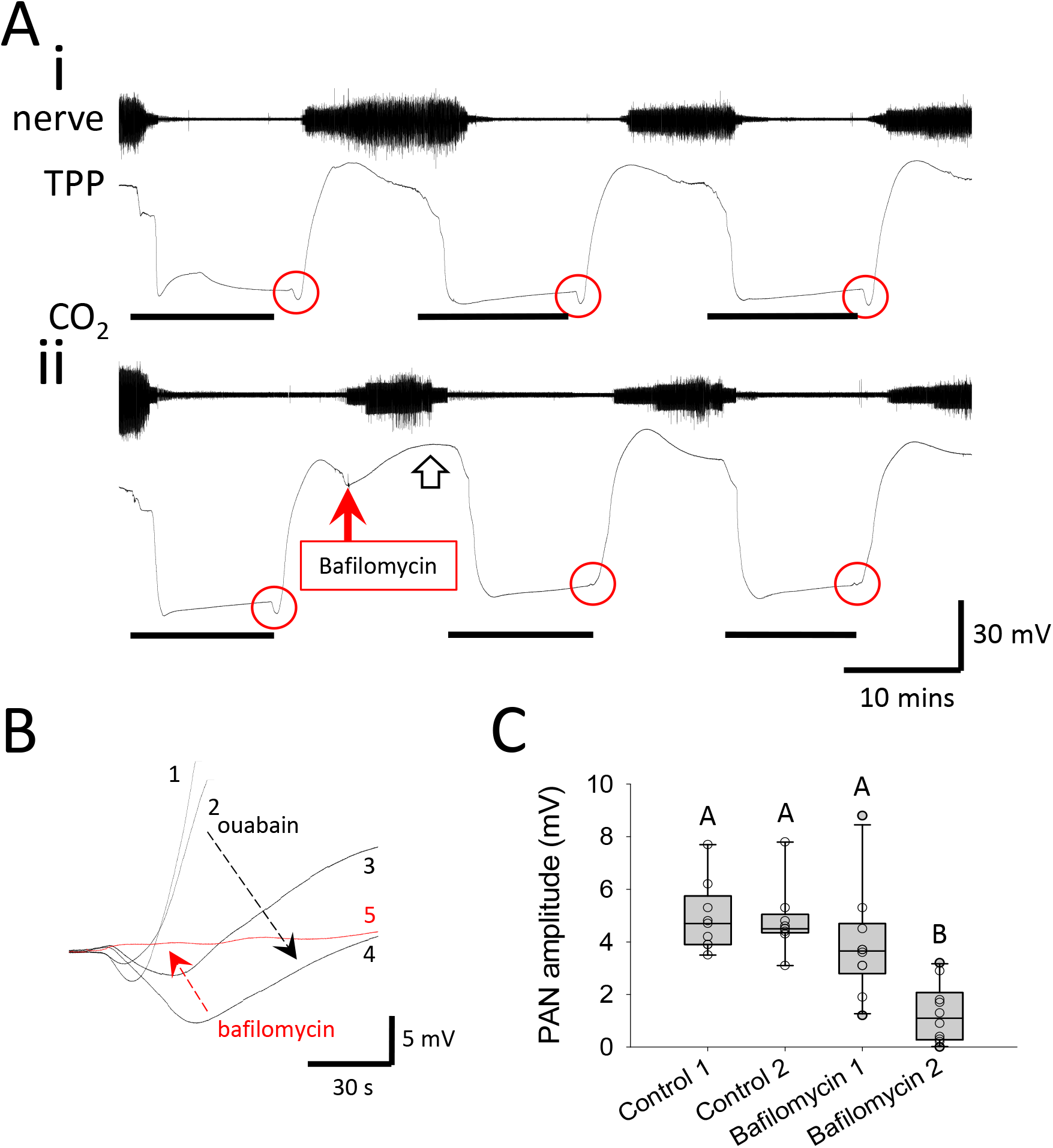
Bafilomycin eradicates the PAN. **A.** Nerve and TPP recordings during 3 repeated 10 min CO_2_ anoxias. **i.** Control preparation **ii.** Experimental preparation in which 10 µM bafilomycin was added to the saline between the first and second anoxia (solid arrow). Note that bafilomycin application was followed by a positive DC shift of TPP (open arrow) and that the PAN (circled) was eradicated after bafilomycin treatment. **B.** Overlaid traces of the PAN with repetitive 10 min N_2_ anoxias at different times after treatment with ouabain and then bafilomycin. Trace 1 – pre-ouabain; trace 2 – 18 mins after 10 mM ouabain; trace 3 – 34 mins after ouabain; trace 4 – 45 mins after ouabain; trace 5 – 68 mins after ouabain and 17 mins after bafilomycin. Note that ouabain accentuated the PAN (black arrow) and bafilomycin subsequently eradicated it (red arrow). **C.** PAN amplitude for first and second anoxias in control experiments and experiments in which bafilomycin had been applied between the first and second anoxias. Box plots indicate the median, 25^th^ and 75^th^ percentiles with whiskers to the 10^th^ and 90^th^ percentiles. Individual data points plotted as open symbols. Statistically significant differences indicated with different letters.

To determine whether bafilomycin affected the response to anoxia, we compared the percentage change in the timing of failure and recovery of the second anoxia compared with the first anoxia (**Fig. 10**). In control preparations, the time to failure was generally longer for a repeat anoxia compared with the first whereas after bafilomycin the time to rhythm failure was shorter and there was minimal change in the time to SD. The change in the time to rhythm failure was −0.6 (−0.8 – 22.9) % in controls and −8.5 (−12.0 – 3.8) % after bafilomycin (Kruskal-Wallis ANOVA: P = 0.08; after transforming the data [ln(x+17)] Student’s t-test P = 0.03) (**Fig. 10A**). The change in the time to SD was 31.5 ± 28.3 % in controls and 3.9 ± 17.3 % after bafilomycin (Student’s t-test: P = 0.02) (**Fig. 10B**). The change in the recovery of the TPP measured as the time from air on to the time at half-amplitude was −7.4 ± 10.7 % in controls and −19.4 ± 2.9 % after bafilomycin (Student’s t-test: P = 0.02) (**Fig. 10C**). Despite an earlier recovery of the TPP after bafilomycin, it took longer for the rhythm to recover: −3.9 ± 13.8 % in controls and 30.7 ± 33.6 % after bafilomycin (Welch’s t-test for unequal variances; P = 0.02) (**Fig. 10D**).

**Figure 10.**
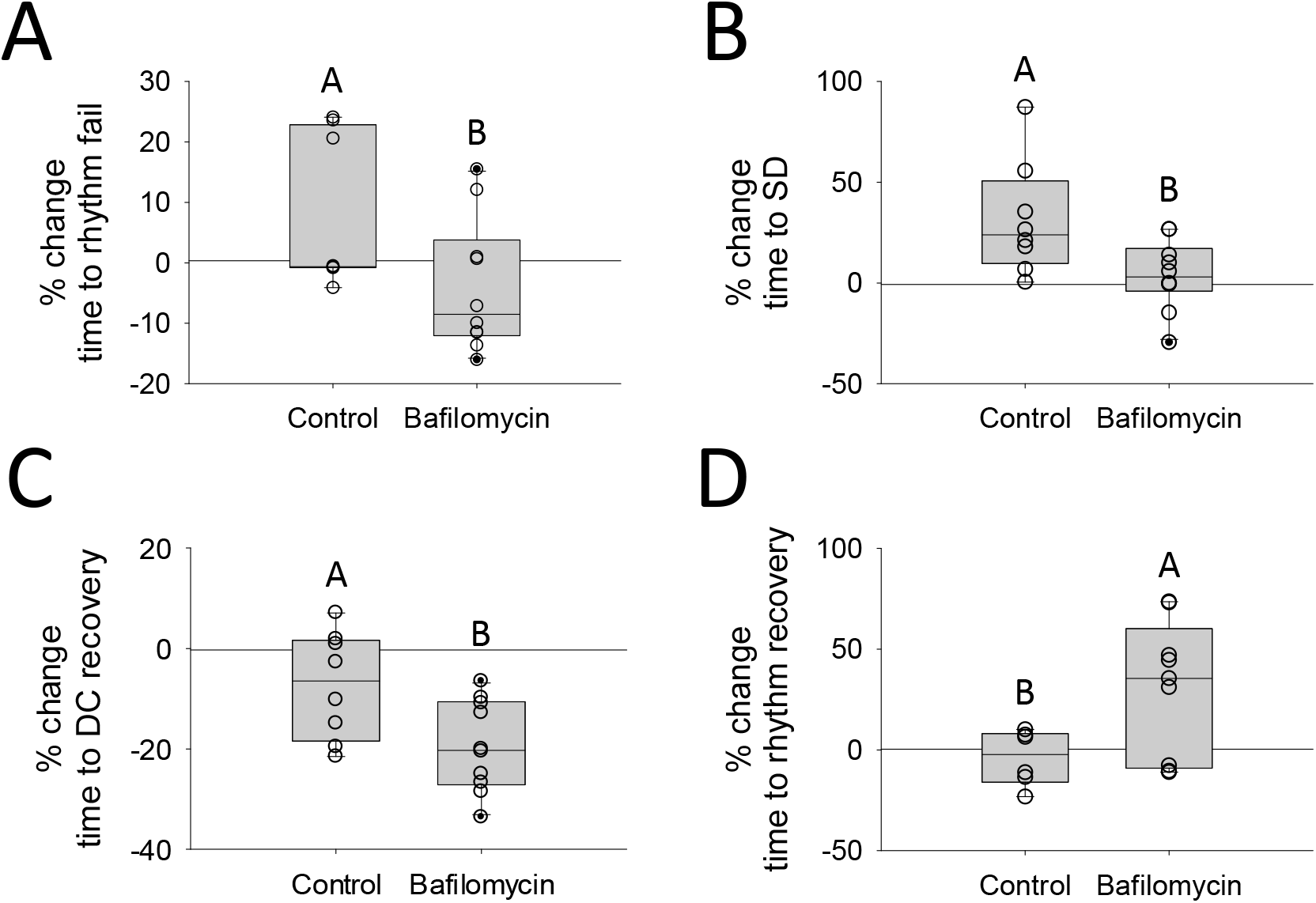
Bafilomycin hastens anoxic coma and delays recovery. Comparison of the time course of failure and recovery with two anoxias in control preparations and before and after 10 µM bafilomycin. Percent changes in: **A.** Time to failure of the ventilatory rhythm. **B.** Time to SD onset. **C.** Time to recovery of the TPP after return to air. **D.** Time to recovery of the ventilatory rhythm. Box plots indicate the median, 25^th^ and 75^th^ percentiles with whiskers to the 10^th^ and 90^th^ percentiles. Individual data points plotted as open symbols. Statistically significant differences indicated with different letters.

#### Summary

Bafilomycin caused a long-term positive DC shift of TPP and eradicated the PAN, consistent with inhibition of the electrogenic VA. In addition, bafilomycin hastened the onset of rhythm failure and coma and delayed the onset of rhythm recovery. Next, we used intracellular recordings to compare the effects of N_2_ and CO_2_ on excitable cells.

### Intracellular Recording

Recording intracellularly from muscle fibres during the onset of anoxia is difficult because of the rapid muscle twitching and contractions prior to coma. Nevertheless, we managed to record successfully in 6 locusts (2 male and 4 female), which were given 7 N_2_ anoxia and 6 CO_2_ anoxia treatments. Recordings were taken from the posterior rotator of the mesothoracic coxa (muscle 93 of (Snodgrass, 1929)) because of its convenient location in this preparation, originating on the second spina between the connectives and immediately anterior to the metathoracic ganglion. Resting membrane potential was −45.8 ± 10.7 mV, which generally hyperpolarized at gas onset prior to the depolarization and burst of activity associate with entry to coma (**Fig. 11**). The extent of the depolarization was not affected by the gas (V_m_ during coma: N_2_ = −25.8 ± 4.5 mV; CO2 = −26.2 ± 5.3 mV; Student’s t-test P = 0.9). However, the latency from anoxia to the beginning of the burst was considerably shorter with CO_2_ anoxia (N_2_ = 2.7 (2.2-3.8) mins; CO_2_ = 0.45 (0.2-0.6) mins; Kruskal-Wallis: P = 0.004). Not surprisingly, there was no change in muscle V_m_ at the time of SD in the ganglion.

**Figure 11.**
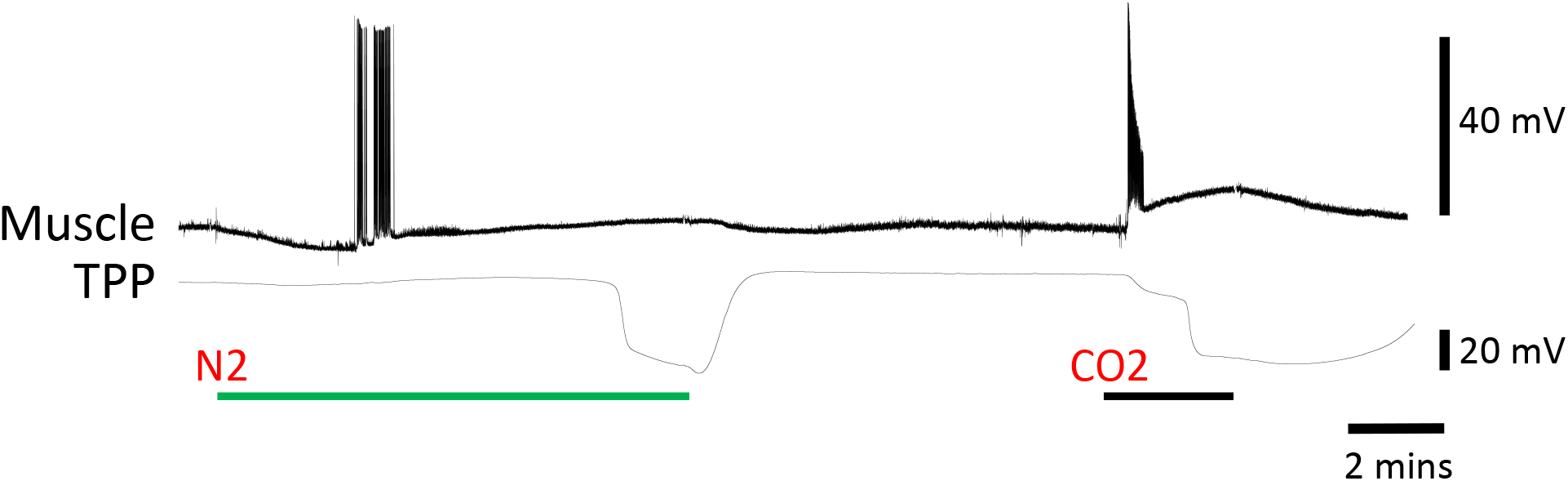
Muscle cells depolarize prior to SD. Recordings from a fibre of the posterior rotator of the mesothoracic coxa and the TPP during 1-minute comas induced by N_2_ and then by CO_2_.

We recorded intracellularly from neurons in 12 locusts (10 male and 2 female), which were given 14 N_2_ anoxias and 11 CO_2_ anoxias. The recordings were taken from the neuropil segments of wing muscle motoneurons, which have extensive arborizations located in a dorsal layer just under the ganglion sheath. These neurons receive a constant barrage of postsynaptic potentials at “rest” and have large, overshooting action potentials. Membrane potential (V_m_) was derived from the intracellular recording relative to ground (V_i_) minus the extracellular recording relative to ground (V_o_ = TPP) (**Fig. 12A**). Initial membrane potential (V_m_) was −69.6 ± 6.5 mV and TPP (V_o_) was 12.7 ± 6.0 mV. Gas onset caused a transient hyperpolarization followed by neuronal activation, which was initially patterned, occasionally with a flight-like rhythm, before turning tonic as the strength of synaptic interactions waned (**Fig. 12B**). During the coma (after SD onset), there was no effect of the nature of the gas on V_m_ (N_2_ = −3.9 ± 6.0 mV; CO_2_ = −7.2 ± 7.6; Student’s t-test: P = 0.25) or on TPP (N_2_ = −35.5 ± 9.4 mV; CO_2_ = −31.6 ± 10.4 mV; Student’s t-test: P = 0.35). Similarly, after recovery there was no effect of the gas on V_m_ (N_2_ = - 71.6 ± 8.6 mV; CO_2_ = −70.9 ± 9.0 mV; Student’s t-test: P = 0.86) or on TPP (N_2_ = 13 ± 5.8 mV; CO_2_ = 14.7 ± 9.7 mV; Student’s t-test: P = 0.6). The primary difference associated with the nature of the gas was that with CO_2_ the depolarization prior to SD was faster and the repolarization on recovery was slower (**Fig. 12C**). Neurons depolarized in two stages, the second stage being simultaneous with the negative DC shift of the TPP. With N_2_, the first stage was relatively gradual, with little change to TPP. Whereas, with CO_2_, the first depolarization was relatively abrupt and V_m_ stepped to −46.6 ± 6.1 mV (∼25 mV depolarization), while TPP stepped to 3.9 ± 8.6 mV (∼10 mV negative DC shift).

**Figure 12.**
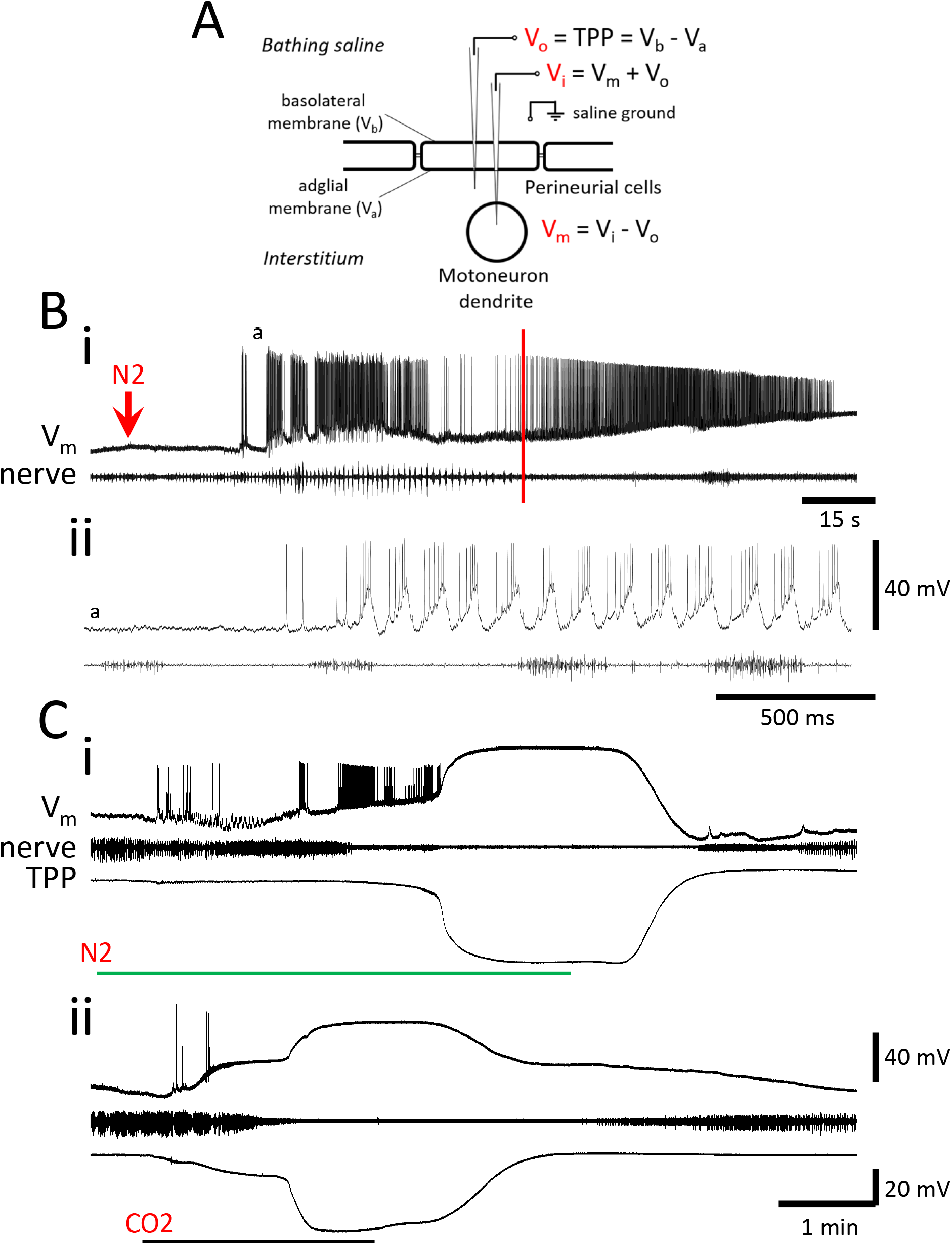
Motoneurons depolarize in two stages. **A.** The recording arrangement. The membrane potential of the motoneuron dendrite (V_m_) was derived from the intracellular potential (V_i_) minus the extracellular potential (V_o_), both recorded relative to the saline ground. V_o_ is the transperineurial potential (TPP) generated by the potentials across the basolateral (V_b_) an adglial (V_a_) membranes of perineurial cells. **B. i.** Intracellular recording from a wing elevator muscle motoneuron and a median nerve at the onset of N_2_ delivery (arrow), prior to SD. The red vertical line indicates the time at which motor patterning fails in the median nerve and motoneuronal firing becomes tonic. **ii.** Expanded portion (a) of the traces in **i**. Note the flight-like bursting activity of the motoneuron and the increasing burst strength of the ventilatory rhythm. **C.** Intracellular recordings from a wing muscle motoneuron, a median nerve and the TPP during **i.** N_2_ anoxia and **ii.** CO_2_ anoxia showing two stages of depolarization. **i** and **ii** are from different preparations.

We wanted to confirm that SD was associated with a membrane conductance decrease of neuronal membranes by recording the voltage response to constant current pulses (1-5 nA). However, this was complicated by the fact that the input resistance across the sheath changed markedly during onset and recovery of the anoxic coma. Normally, for intracellular measurement of neuronal input resistance the effect of the V_m_ voltage drop across the electrode resistance (variable in different experiments) is cancelled by balancing a bridge circuit of the amplifier. This is acceptable for V_i_ measures relative to ground if V_o_ does not change. However, in these experiments, V_o_ and the input resistance of the sheath changed dramatically at SD onset and this contaminates the intracellular V_i_ recording.

Prior to anoxia the R_in_ of the sheath was 0.14 ± 0.04 MΩ (n = 3). In 10 male locusts, we measured the changes in input resistance of the sheath using constant current pulses (**Fig. 13A**). At about the time that patterning of activity in the nerve recording ceased, the sheath R_in_ started to gradually increase until there was a more abrupt increase at SD onset. During the coma, sheath R_in_ was constant and 3.7 ± 1.8 times the initial value. This gradually returned to 0.75 ± 0.47 times the initial value after recovery. The increased R_in_ during SD was different from initial and recovered values (One-way RM ANOVA followed by Bonferroni posthoc comparisons: P < 0.001). There was no difference between initial and recovered values (Bonferroni: P = 1.0).

**Figure 13.**
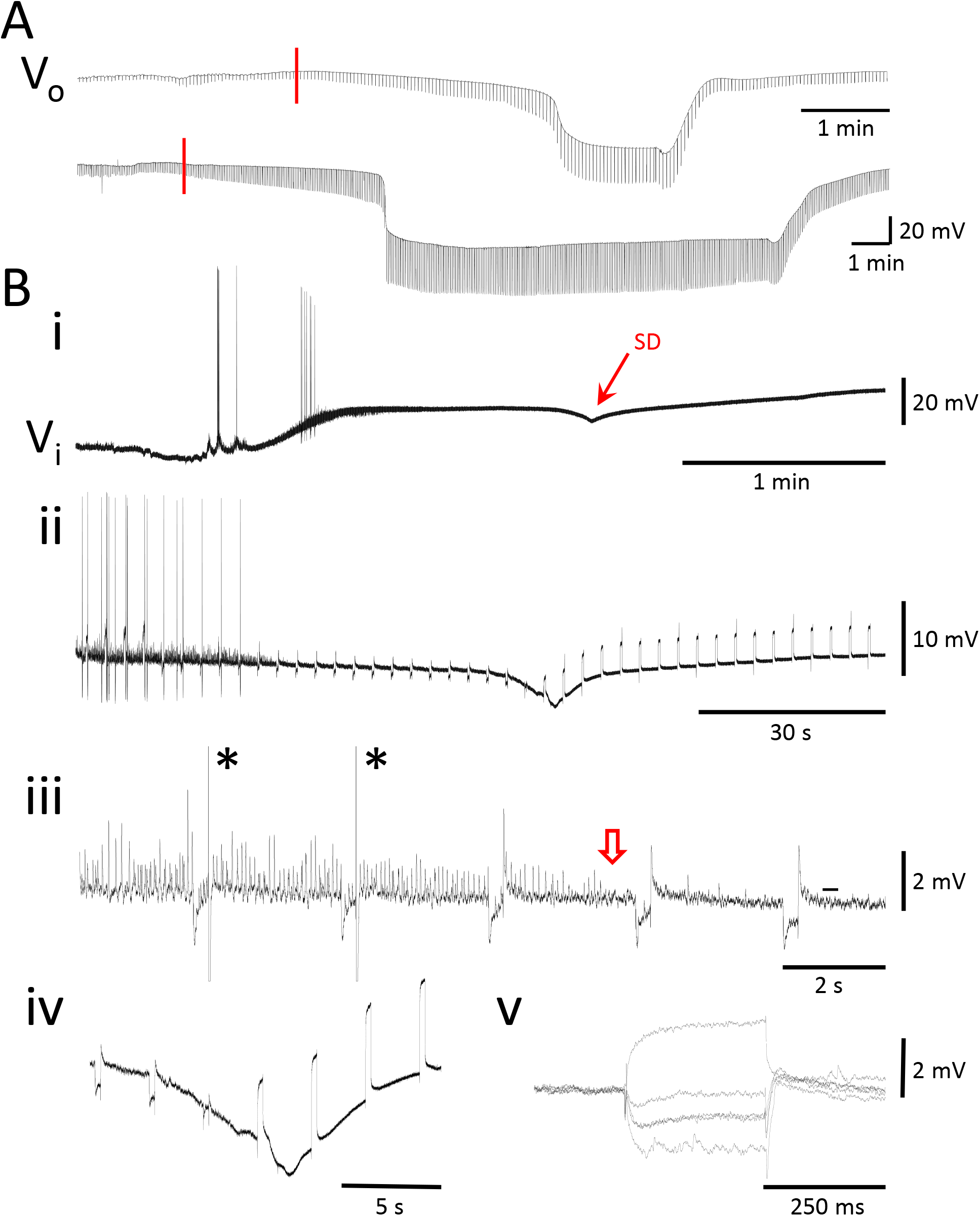
Input resistance recordings. **A**. Constant current pulses delivered to TPP (V_o_) recordings for a 1-minute (upper) and a 10-minute (lower) N_2_ anoxia. Traces start at the onset of N_2_. The red lines indicate when motor patterning fails in a median nerve (not shown). Note the increases in sheath Rin, indicated by the increased voltage deflections, that remains constant during the comas. **B. i.** Intracellular recording relative to ground (V_i_) from a wing muscle motoneuron at the onset of CO_2_-induced SD. Same recording as shown in Fig. 11Cii, which is the derived V_m_ after subtracting V_o_ (TPP). Note the notch in the recording at the time of SD onset. **ii.** A different preparation with constant current pulses. **iii.** Expansion of **ii** indicating action potentials generated by post-inhibitory rebound (asterisks) and the failure of synaptic transmission (open arrow). **iv.** Expansion of **ii** indicating the change in the voltage responses to constant current pulses at the onset of SD. **v.** Overlaid voltage responses to constant current pulses prior to anoxia (lowest, with synaptic potentials) and around SD onset (the first 4 pulses in **iv**). Note the decrease in amplitude indicating decreased Rin of the neuron and the abrupt amplitude increase indicating loss of bridge balance.

In some intracellular recordings it was possible to examine the voltage response to current pulses and discount the immediate voltage shift cause by an unbalanced bridge and measure only the exponential change in voltage that would be expected of the voltage drop across a membrane capacitance. Doing this, in 7 male locusts with 8 N_2_ anoxias and 6 CO_2_ anoxias, neuronal R_in_ was initially 2.3 ± 1.1 MΩ and there was no significant change until SD onset. At SD onset, R_in_ decreased to 0.3 ± 0.5 MΩ, returning to 2.0 ± 0.8 MΩ on recovery; there was no effect of the gas (Two-way RM ANOVA: P_gas_ = 0.70; P_timing_ < 0.001; P_gas x timing_ = 0.82). In the V_i_ recordings of all CO_2_ anoxias and some N_2_ anoxias, there was a clear negative notch in the voltage at the onset of SD (**Fig. 13Bi,ii**). This was simultaneous with the decrease in neuronal membrane resistance and the increase in sheath input resistance (**Fig. 13Biii,iv,v**).

#### Summary

Muscle fibres depolarized during anoxia. The nature of the gas did not affect the extent of the depolarization, but CO_2_ had a more rapid onset and slower recovery. Neurons depolarized in two stages during anoxia. CO_2_ caused a relatively rapid initial depolarization prior to SD. There was an abrupt conductance increase in neurons that occurred at SD onset and was not affected by the nature of the gas. Recovery was slower with CO_2_ anoxia.

## Discussion

Locusts recover the ability to stand after 6 hours in a N_2_-induced coma (Wu et al., 2002). However, recovery is incomplete with muscle tissue damage after 4 hours of anoxia (Ravn et al., 2019) and they have been known to die without feeding 3-5 days after only 2 h of anoxia (Michel and Wegener, 1982). In adult *Drosophila,* the ability to withstand anoxia is related to the maintenance of hypometabolism and tolerance of ionic variability (Campbell et al., 2018; Campbell et al., 2019). We were interested in the contribution of the CNS to hypometabolism and in the effects of different methods of inducing anoxia prior to any permanent injury. Thus, we used anoxia durations that result in apparently complete recovery (≤ 30 mins). We found that at the level of the CNS there was little difference between the effects of water immersion and those of 100% N_2_ treatment. However, although intact locusts recovered faster from a CO_2_ anoxia than from N_2_ or H_2_O, the effects of CO_2_ in the CNS, which were more rapid and intense at onset, took longer to dissipate. The slower recovery of CNS operation with CO_2_ was associated with a slow recovery from a much more pronounced interstitial acidosis and a greater activation of a V-ATPase. SD, when it occurred, was not obviously affected by the nature of the gas, suggesting that its mechanisms were unchanged. At SD onset, the adglial membrane of perineurial glial cells depolarized before deeper neuronal membranes, indicating that anoxic SD propagates from an event initiated at the perineurial layer of the BBB. This intriguing result provides a novel perspective on SD mechanisms considering the density of mitochondria packed into perineurial glial cells surrounding the ganglia (Smith and Shipley, 1990).

### Motor patterning

The first sign of hypoxia was an increase in the frequency of abdominal pumping movements. Nerve recording of the underlying motor pattern showed that this occurred immediately, before any change in pH_o_ or [K^+^]_o_ was recorded. The central pattern generator for ventilation is located in the metathoracic ganglion (Bustami and Hustert, 2000) where detection of O_2_ and CO_2_ levels is likely a widespread property of neural tissue rather than being located in specific regions (Bustami et al., 2002; Talal et al., 2019). A recent model of O_2_ chemoreception in glomus cells of the mammalian carotid body proposes that acute detection of reduced O_2_ is a function of mitochondrial complex IV with subsequent mitochondrial signalling via NADH and reactive oxygen species to membrane ion channels (Ortega-Saenz and Lopez-Barneo, 2020; Ortega-Saenz et al., 2020). Hence, to a greater or lesser extent, neurons will respond to metabolic perturbation of mitochondria; indeed this is recognized as a mechanism for the homeostatic regulation of neuron excitability (Ruggiero et al., 2021). After recovery, the frequency of the ventilatory motor pattern was greatly reduced by N_2_ and CO_2_ anoxia but not by water immersion. Mammalian ventilatory reflexes can be facilitated by the energy sensor, AMPK, which is activated by an increasing AMP:ATP ratio (Evans, 2019; Evans and Hardie, 2020), also implicating the involvement of mitochondrial operation. In locusts, the ventilatory motor pattern changes induced by mitochondrial inhibition using sodium azide (chemical “anoxia”) are mimicked by AMPK activation using AICAR, although in the latter case there is no SD (Rodgers-Garlick et al., 2011). Given the difference in the motor pattern changes after recovery, our current results suggest that water immersion was less metabolically stressful than N_2_ or CO_2_ exposure. This may have been a consequence of residual O_2_ in the tracheae at coma onset.

The second stage in the behavioural response to increasing hypoxia was a transition from vigorous ventilation to immobility. In nerve and neuron recordings, this was associated with an abrupt transition from patterned neural activity to tonic firing. Initially we ascribed this to a failure of synaptic transmission, however synaptic potentials were recorded in neurons up to the point when excitability failed. Moderate excitation increased ventilatory activity and could activate latent circuitry (e.g., to release flight-like motor patterns). The failure of motor patterning may have been due to a reduction of synaptic potential amplitude below a threshold required for circuit operation. Alternatively, the excitatory effects of extreme hypoxia may have generated spike frequencies that preclude patterning i.e., by rendering firing neurons unresponsive to inhibitory inputs. This could be resolved by monitoring identified synapses (e.g. from the wing hinge stretch receptor to flight interneurons (Gee and Robertson, 1994)) before, during and after anoxia. The recovery of motor patterning is clearly dependent on the recovery of synaptic transmission in neuronal circuits. In mammalian preparations, the recovery of synaptic transmission after SD is delayed by an accumulation of adenosine, which presynaptically inhibits transmitter release (Lindquist and Shuttleworth, 2012; Lindquist and Shuttleworth, 2017). Adenosine also delays functional recovery after anoxic comas in locusts but does not affect the timing of SD (Van Dusen et al., 2020a).

The effects of anoxia on circuit function are distinct from anoxic SD. Whereas SD arrests all neural function, metabolic stress can affect motor patterning in the absence of SD. Moreover, the timing of motor pattern failure and recovery can be modified independently from the temporal characteristics of SD. This suggests that the mechanisms of SD are independent from the specific mechanisms underlying action potential generation and synaptic transmission.

### Sex

After anoxia, male locusts recover CNS operation more slowly than females (Hou et al., 2014; Robertson et al., 2019; Van Dusen et al., 2020a). Our results confirm that males recover ventilation more slowly than females after N_2_ anoxia. In the Australian Plague Locust, this sex difference develops during maturation of adults in the gregarious phase, at the time when adults start mating, suggesting that it is associated with increased CNS metabolic rate of males competing for mates in a crowded environment (Robertson et al., 2019). The fact that females recovered the ability to stand after CO_2_ anoxia more slowly than males may reflect prolonged effects of CO_2_ anesthesia on a larger muscle mass in females. The timing of behavioural recovery after anoxia is positively correlated with the duration of the coma due to the build-up of metabolites (e.g., adenosine; see above) that take time to clear (Lighton and Schilman, 2007; Weyel and Wegener, 1996). Moreover, recovery from a prior anoxia, a treatment known to reduce neural performance and whole animal metabolic rate via activation of AMPK (Money et al., 2014), reduces the time to recovery from a subsequent anoxia (Robertson et al., 2019). Thus, energy metabolism during the coma will have an impact on the timing of recovery. It is important to note that recovery of the CNS, while obviously permissive for recovery of the whole animal, may be differentially modulated by neural conditions and/or neuromodulators that could have a minimal or different impact on the intact locust. Neural energetics in insects are under tissue-specific neuromodulatory control in ways that support age-dependent or phase-dependent behaviours (Rittschof and Schirmeier, 2018; Rittschof et al., 2019).

An additional sex-difference was that, after 10 mins of N_2_ coma, the amplitude of the PAN of males was twice that of females. We found that the PAN was associated with a transient decrease in pH_o_ and could be eradicated by pretreatment with bafilomycin, an inhibitor of VA, a proton pump. We attribute the PAN to the electrogenic effect of VA and its negative quality indicates that protons were being cleared from the interstitium into the hemolymph (bathing saline). The fact that the amplitude of the PAN was strongly and positively correlated with the duration of the coma suggests that it reflects the build-up of protons derived from anaerobic metabolic activity. This would parallel the build-up of the anaerobic end product, lactate, which has been described for hemolymph of *Acheta domesticus* (Woodring et al., 1978), whole animals and flight muscle of *Schistocerca gregaria* (Hochachka et al., 1993), pupae of *Manduca sexta* (Woods and Lane, 2016) and adults and larvae of *Drosophila melanogaster* (Campbell et al., 2019). Thus, we propose that, compared to females, male locusts have higher levels of anaerobic glycolysis in the CNS during anoxia, which results in a slower CNS recovery (i.e., slower recovery of ventilatory rhythm generation).

### pH

A major contributor to pH_o_ with CO_2_ anoxia is clearly the gas delivery generating carbonic acid and causing an immediate decrease of ∼1 pH unit. Nevertheless, with both CO_2_ and N_2_ anoxia, a slower interstitial acidification was likely due to hyperactivity (Rasmussen et al., 2020) and anaerobic glycolysis with the production of protons. Restoration of pH_o_ was slower than restoration of [K^+^]_o_ and was interrupted by transient acidification associated with the return of neural activity. Activity causes extracellular acidification in neural tissue (Chesler, 2003; Magnotta et al., 2012; Xiong and Stringer, 2000) some of which is due to protons released by synaptic activity (Chiacchiaretta et al., 2017); synaptic vesicles are acidified by VA to facilitate transmitter loading (Mellman et al., 1986). In rat cortex, spreading depression and cerebral ischemia causes pH_o_ to drop from 7.33 to 6.97 and 6.75 units (respectively); restoration of pH_o_ after spreading depression parallels restoration of lactate levels (Mutch and Hansen, 1984).

In our experiments the large decrease of pH_o_ induced by CO_2_ is unlikely to have caused the increased ventilatory frequency, which occurred immediately with N_2_ anoxia without any pH_o_ change. Acidification of the hemolymph does not increase ventilation rate in grasshoppers (Gulinson and Harrison, 1996; Krolikowski and Harrison, 1996). In our semi-intact preparations, up to an hour of exposure to pH 3.5 saline had no effect on ventilatory rhythm frequency (n = 3; RMR unpublished observations). However, the ganglion sheath is a very effective barrier to protons and these treatments may not have changed pH_o_. Interstitial acidification generally reduces excitability of neurons and synaptic transmission (Chesler, 2003; Rasmussen et al., 2020; Tombaugh and Somjen, 1996), as we noted at the start of anoxia in intracellular neuron and muscle fibre recordings, and it is unlikely to be responsible for the hyperexcitability prior to coma onset. We found that inhibition of VA with bafilomycin, which would have slowed restoration of pH_o_, shortened the time to rhythm failure and entry to coma and increased the time taken to restore motor pattern generation. This underlines the importance of pH homeostasis for proper CNS operation. Nevertheless, we do not know the effects of interstitial acidification on neural mechanisms in our preparation and it would be interesting to directly manipulate pH_o_ by injection across the ganglion sheath.

### Transperineurial potential

The TPP is a convenient indicator of the occurrence and timing of SD. It depends on basolateral and adglial membrane potentials of the perineurial cells of the BBB (Schofield and Treherne, 1984). In turn, these depend on many different parameters that can vary independently of each other (ion concentrations of hemolymph, interstitium and cytosol; ion conductances of the two membranes; electrogenic activities of energy-dependent ion pumps). Thus, it may be misleading to focus on TPP dynamics as an indicator of failure and recovery of the CNS. What is functionally important in the CNS is the failure and recovery of synaptic transmission and action potential generation. The TPP has no intrinsic functional relevance but measuring the TPP can provide information about mechanisms that might underlie the failure and recovery of neural operation. Interpretation of TPP dynamics is complicated. The fact that it is negative during SD may be of no functional relevance apart from what it indicates about the adglial membrane potential depolarizing close to zero while the basolateral membrane maintains a negative membrane potential, indicating continuing integrity of the BBB.

The amplitude of the PAN of the TPP correlates positively with TPP recovery time because a larger initial negative excursion will necessarily delay the return of TPP to starting values, assuming the restoration rate is the same. Also, reduction of the PAN using bafilomycin to inhibit the VA shortened TPP recovery time, but it increased the time to recovery of ventilatory motor patterning (synaptic transmission). An interpretation is that the timing and slope of TPP recovery are determined by overlapping phenomena that are all restorative but push the TPP in different directions (e.g., negative shift for VA and positive shift for NKA). The TPP provides a good general indicator of when SD starts and stops but the details of its trajectory, including amplitude, need careful interpretation.

Intracellular recording with sharp electrodes relative to the bathing medium at ground (V_i_) is complicated by the changes of TPP induced by anoxia and SD. Under normoxia, the intracellular electrode can be zero-ed in the interstitium, after penetrating the sheath, and the resulting recording with be a faithful representation of V_m_ because small activity-dependent variation in TPP (V_o_) has negligible effect. The large changes of TPP during SD can not be ignored. This is complicated by the fact the TPP depends on both the basolateral and adglial membrane potentials, which can change independently. At SD, adglial and neuronal membranes both depolarize close to zero and the changes to V_m_ and V_o_ (TPP) will be equal and opposite, cancelling each other out and resulting in almost no change in V_i_. We did, however, notice a notch in the V_i_ recording at SD onset. This was always negative and more pronounced with CO_2_ anoxia. We interpret this as being due to a slight mismatch in the timing of depolarization of the adglial and neuronal membranes. The fact that the notch was always negative indicates that the adglial membrane depolarizes first (negative shift of V_o_) followed several seconds later by the depolarization of the neuronal membrane (positive shift of V_m_). The fact that it was more noticeable with CO_2_ anoxia is because this initially generates a more abrupt and larger neuronal depolarization providing a depolarization that enhances the appearance of the negative shift of V_o._ We propose that the wave of SD is initiated at the BBB, depolarizing glial membranes first, and propagates both laterally, generating latency differences of the negative DC shift, and more deeply, depolarizing neuronal membranes and generating the notch in the V_i_ recordings.

Constant current pulses delivered during a V_o_ recording showed that the electrical pathway from the interstitium to the bathing medium increased in resistance. This started gradually from around the time that motor patterning failed and increased more abruptly at SD onset, remaining steady during the coma, and returning to starting values with the return to normoxia. This could have been caused by conductance changes across or between the perineurial cells (not the adglial membrane, which is depolarized, presumably because of an increased conductance). An alternative explanation is that cell swelling (Spong et al., 2015), which occurs with SD and is the basis for optical recording of SD progression (Anderson and Andrew, 2002), compressed the extracellular pathway, increasing its resistance. At present, we suspect that both mechanisms have a role but cannot distinguish their relative importance. Nonetheless, these findings illustrate that the ganglion sheath remains an effective barrier, indeed increasing its efficacy, to the free flow of ions during SD.

Another complication of the V_i_ recording is evident when using the delivery of constant current pulses to characterize input resistance of neurons. The resistance changes in the extracellular pathway noted above contaminated the intracellular recordings and prevented stable balancing of the amplifier bridge circuit. In spite of that, we are confident that SD onset was associated with an abrupt decrease in the resistance of neuronal membranes as has been previously described in mammalian brain slices (Czéh et al., 1993) and for azide-induced SD in the locust metathoracic ganglion (Armstrong et al., 2009).

## Conclusions

Our goal was to understand the mechanisms underlying the differences between the effects of water immersion or gas (N_2_ or CO_2_) exposure for inducing anoxia in locusts. At the level of the CNS there was little difference between water immersion and N_2_ exposure and whole animal differences can be attributed to characteristics of the tracheal system. Thus, the reservoir of air in the tracheae at the time of immersion prolongs the time to enter a coma. Recovery from water immersion may have been hindered by residual water collected in the spiracles.

Whole animal recovery was quickest after CO_2_ anoxia. Given that this was not evident in the CNS, where recovery from CO_2_ was slowest, the difference must be due to peripheral mechanisms. An explanation is provided by the anesthetic effects of CO_2_ inhibiting neuromuscular transmission by decreasing the sensitivity of glutamate receptors (Badre et al., 2005). Rapid shutdown of neuromuscular transmission would prevent depletion of transmitter at neuromuscular junctions allowing a more rapid functional recovery. There may be other differences within muscle fibres that are protected by early paralysis to promote recovery of muscle strength.

The effect of CO_2_ on the whole animal resembles in some fashion the effect of muscle relaxants in the context of electroconvulsive therapy for humans; although there is sudden and uncoordinated hyperactivity of neurons, this is not evident in the behaviour of the animal. All our results show that, compared with N_2_, CO_2_ causes an immediate and more extreme hyperexcitability and greater interstitial acidification from which it takes longer to recover. There was no obvious difference in the characteristics of SD itself and it is pertinent that although CO_2_ hastened the onset of SD it did not alter the propagation speed. It is tempting to ascribe the differences in excitability and SD onset to the substantial acidification cause by CO_2,_ but we do not yet have direct evidence to make that connection. Also, it is worth considering that the CO_2_ treatment is more severe in the sense that the concentration changes suddenly from 0.04% in air to 100%, in addition to 0 % O_2_, whereas with N_2_ the change is primarily in the loss of O_2_. Arguably, it is to be expected that the consequences, although essentially the same, would be more severe with CO_2_. The mechanisms underlying anoxic SD in the CNS were not noticeably different with the different methods of anoxia. Future research will focus on how events at the perineurial glial layer trigger SD that propagates deeper into the neuropil.

## Abbreviations

AICAR: 5-aminoimidazole-4-carboxamide ribonucleoside
AMPK: AMP-activated protein kinase
BBB: blood brain barrier
EMG: electromyographic
MTG: metathoracic ganglion
NKA: Na^+^/K^+^-ATPase
PAN: postanoxic negativity
pH_o_: interstitial pH of the ganglion
*P*_O2_: partial pressure of oxygen
SD: spreading depolarization
TPP: transperineurial potential
V_i_: intracellular potential relative to the bathing saline
V_m_: membrane potential = (V_i_-V_o_)
V_o_: extracellular potential relative to the bathing saline
VA: vacuolar-type (V)-ATPase

## Acknowledgements

We thank Laura Dwyer for collecting the data in Figure 1 and Mike O’Donnell for suggestions about ion-sensitive electrode recording and the V-ATPase. We also thank Mads Andersen, David Andrew, Heath MacMillan, Chris Moyes, Mike O’Donnell and Yuyang Wang for their comments on a previous version of the manuscript. Funded by a Discovery Grant from the Natural Sciences and Engineering Research Council of Canada.

